# Depolymerizing F-actin accelerates the exit from pluripotency to enhance stem cell-derived islet differentiation

**DOI:** 10.1101/2024.10.21.618465

**Authors:** Nathaniel J. Hogrebe, Mason D. Schmidt, Punn Augsornworawat, Sarah E. Gale, Mira Shunkarova, Jeffrey R. Millman

## Abstract

In this study, we demonstrate that cytoskeletal state at the onset of directed differentiation is critical for the specification of human pluripotent stem cells (hPSCs) to all three germ layers. In particular, a polymerized actin cytoskeleton facilitates directed ectoderm differentiation, while depolymerizing F-actin promotes mesendoderm lineages. Applying this concept to a stem cell-derived islet (SC-islet) differentiation protocol, we show that depolymerizing F-actin with latrunculin A (latA) during the first 24 hours of definitive endoderm formation facilitates rapid exit from pluripotency and alters Activin/Nodal, BMP, JNK-JUN, and WNT pathway signaling dynamics. These signaling changes influence downstream patterning of the gut tube, leading to improved pancreatic progenitor identity and decreased expression of markers associated with other endodermal lineages. Continued differentiation generates islets containing a higher percentage of β cells that exhibit improved maturation, insulin secretion, and ability to reverse hyperglycemia. Furthermore, this latA treatment reduces enterochromaffin cells in the final cell population and corrects differentiations from hPSC lines that otherwise fail to consistently produce pancreatic islets, highlighting the importance of cytoskeletal signaling at the onset of directed differentiation.

## INTRODUCTION

Pancreatic β cells regulate glucose metabolism throughout the body by secreting insulin into the bloodstream, facilitating glucose uptake by many cell types so that it can be utilized for energy production and fat storage.^1–3^ Within the pancreas, β cells are clustered together with other endocrine cell types to form the islets of Langerhans. In type 1 diabetes (T1D), β cells are erroneously targeted by the immune system and selectively eliminated.^4,5^ As a result, T1D patients must inject exogenous insulin in order to restore proper insulin signaling throughout the body, which is required for survival. Replicating the precise insulin secretion kinetics of β cells with insulin injections can be difficult because insulin requirements change dynamically based on factors related to energy metabolism, such as the amount and types of food consumed, the duration and intensity of physical activity, and the levels of circulating stress hormones. Over time, chronically elevated blood glucose levels can lead to severe complications, such as cardiovascular, kidney, and eye disease, emphasizing the importance of keeping blood glucose levels as close to normal as possible.^6^

In an effort to improve blood glucose control and ease the burden of diabetes management, primary β cells from deceased donors have been transplanted into patients with T1D, demonstrating that replacing this lost β cell mass is a safe and effective method to restore glucose homeostasis.^7,8^ However, issues with donor quality, quantity, and immunogenicity have hampered its widespread use. As an alternative, stem cell-derived islets (SC-islets) containing stem cell-derived β cells (SC-β cells) and other pancreatic endocrine cell types could provide an unlimited source of islets for transplantation.^9^ These SC-islets can be derived from a single stem cell source that is extensively characterized, resulting in more reproducible transplant outcomes. Furthermore, these cells can be gene-edited to have desirable characteristics that enhance transplant results, such as decreasing their immunogenicity to circumvent the need for immunosuppressive drugs^10–14^ or making them more resistant to the stresses experienced during transplantation.^15,16^ SC-islets can also be utilized as an effective *in vitro* model to study different aspects of the disease and develop additional therapeutic strategies.^17–21^

To this end, directed differentiation protocols have been developed in recent years to generate SC-islets using a stepwise combination of growth factors and small molecules to drive hPSCs through several intermediate cell types on their way to becoming pancreatic endocrine cells.^22–29^ While there are a number of different iterations and improvements that have been made since the original development of these protocols, all successful SC-islet differentiation methods follow the same basic strategy. Specifically, they attempt to recreate the specific sequence of signaling events that must occur to form the pancreas during embryonic development.^30^ This process begins with the specification of the definitive endoderm germ layer from hPSCs, which is driven by a coordination of Activin/Nodal, WNT, and BMP signaling.^31–33^ This definitive endoderm is then specified as the primitive gut tube, which will eventually form the organs of the gastrointestinal and respiratory tracts. The primitive gut tube is progressively segmented into the foregut, midgut, and hindgut through signaling gradients of FGF, BMP, WNT, and retinoic acid (RA) ligands that are secreted from the splanchnic mesodermal tissue surrounding the gut tube, as well as Hedgehog and Notch signaling originating in the gut tube itself. ^30,31,34,35^ The intensity and timing of these signals induce gene expression patterns that segment the gut tube into the various organ-forming regions. For example, SOX2 expression is restricted to the foregut, while CDX2 expression is localized to the mid- and hindgut.^36^ Furthermore, differential expression of various HOX genes along the gut tube provide a spatial code that helps guide these different regions to develop into their respective organs.^34,37^ Cells of the foregut will subsequently generate organs such as the thyroid, thymus, trachea, lungs, esophagus, stomach, pancreas, and liver, while the midgut forms the small intestine and the hindgut develops into the large intestine. The pancreas arises from the posterior segment of the foregut, where high RA signaling helps induce the expression of PDX1. These multipotent PDX1+ pancreatic progenitors subsequently develop into duct and acinar exocrine cells as well as all the endocrine cell types of the islet.

Throughout embryonic development, individual cell migration and force generation drive whole tissue movements that result in cell organization and organ formation.^38,39^ These forces are generated within each cell predominately by the actin cytoskeleton and its binding partners, such as myosin motor proteins. In order to generate these forces, actin monomers must polymerize to form filamentous actin (F-actin). This cytoskeletal rearrangement process is dynamic, as the cell continuously integrates multiple intrinsic and extrinsic signaling cues to adopt the proper cytoskeletal configuration for its current stage of development.^39^ Importantly, changes in cytoskeletal state not only control the extent of force generation but have also been shown to be critical to a variety of biochemical signaling pathways and cell processes.^40,41^ While there is also emerging evidence for the importance of cytoskeletal dynamics specifically in hPSCs, much is still unknown on how it affects their behavior compared to other cell types.^42^

In this study, we modulated the state of the actin cytoskeleton during the exit of hPSCs from pluripotency when differentiating them to all three germ layers, demonstrating improved lineage specification depending on the treatment and differentiation protocol. Specifically, a depolymerized cytoskeleton promoted mesendoderm lineages in their respective differentiation media, while a polymerized actin cytoskeleton facilitated directed ectoderm differentiation.

Applying these findings specifically to an SC-islet differentiation protocol, we demonstrated that depolymerizing F-actin with the compound latA during the first 24 hours of differentiation to definitive endoderm facilitated a more robust exit from pluripotency and induced a unique pattern of Activin/Nodal, BMP, JNK-JUN, and WNT signaling. These latA-treated definitive endoderm cells were able to subsequently differentiate into SC-islets with a higher percentage of SC-β cells as well as fewer enterochromaffin cells, which are a type of intestinal endocrine cell that is typically generated during these SC-islet differentiation protocols.^43^ Importantly, depolymerizing the actin cytoskeleton with latA during the first 24 hours of differentiation drastically improved multiple components of SC-β cell function by the end of the protocol 5 weeks later, including higher insulin content and increased insulin secretion in response to glucose stimulation. Not only did this latA treatment improve insulin secretion of two stem cell lines that already generated functional SC-islets reliably, but it also rescued two hPSC lines that failed to consistently produce SC-islets. Overall, this work demonstrates the broad importance of cytoskeletal state during the exit of hPSCs from pluripotency as well as highlights how early interventions that refine patterning of the gut tube to better specify pancreatic progenitors can improve SC-islet generation.

## RESULTS

### Actin polymerization state at the onset of differentiation influenced stem cell specification to all three germ layers

Previously, we demonstrated that the polymerization state of the actin cytoskeleton was critical for the conversion of pancreatic progenitors to pancreatic endocrine and exocrine cells.^26^ Specifically, a highly polymerized actin cytoskeleton induced by culture on stiff polystyrene plasticware blocked NEUROG3-induced endocrine formation, keeping the cells in a pancreatic progenitor state despite the presence of endocrine-inducing factors. Depolymerizing F-actin at the onset of endocrine induction with the compound latA, however, facilitated the expression of NEUROG3 and the initiation of the endocrine program even when the cells continued to be grown on the stiff tissue culture polystyrene substrate.

Since cytoskeleton signaling is important to tissue morphogenesis throughout development and is highly context dependent,^38,39^ we wanted to investigate if cytoskeletal state was also important to other stages of the *in vitro* directed differentiation of hPSCs to SC-islets. While chemically modulating the cytoskeleton at various other stages most often negatively influenced differentiation outcomes, we observed a more diverse response to cytoskeletal manipulation during definitive endoderm formation (Fig. 1A-B, Supplemental Fig. 1A). In particular, adding compounds that depolymerized F-actin during the first 24 hours of stage 1 (Supplemental Fig. 1A) resulted in SC-islets that secreted more insulin in response to glucose stimulation by the end of the 5-week protocol (Fig. 1C). In contrast, compounds that increased actin polymerization during this initial 24-hour period generated non-endocrine cell types (Fig. 1B, Supplemental Fig. 1C-D) and decreased insulin secretion of the final cell clusters (Fig. 1C).

**Figure 1:**
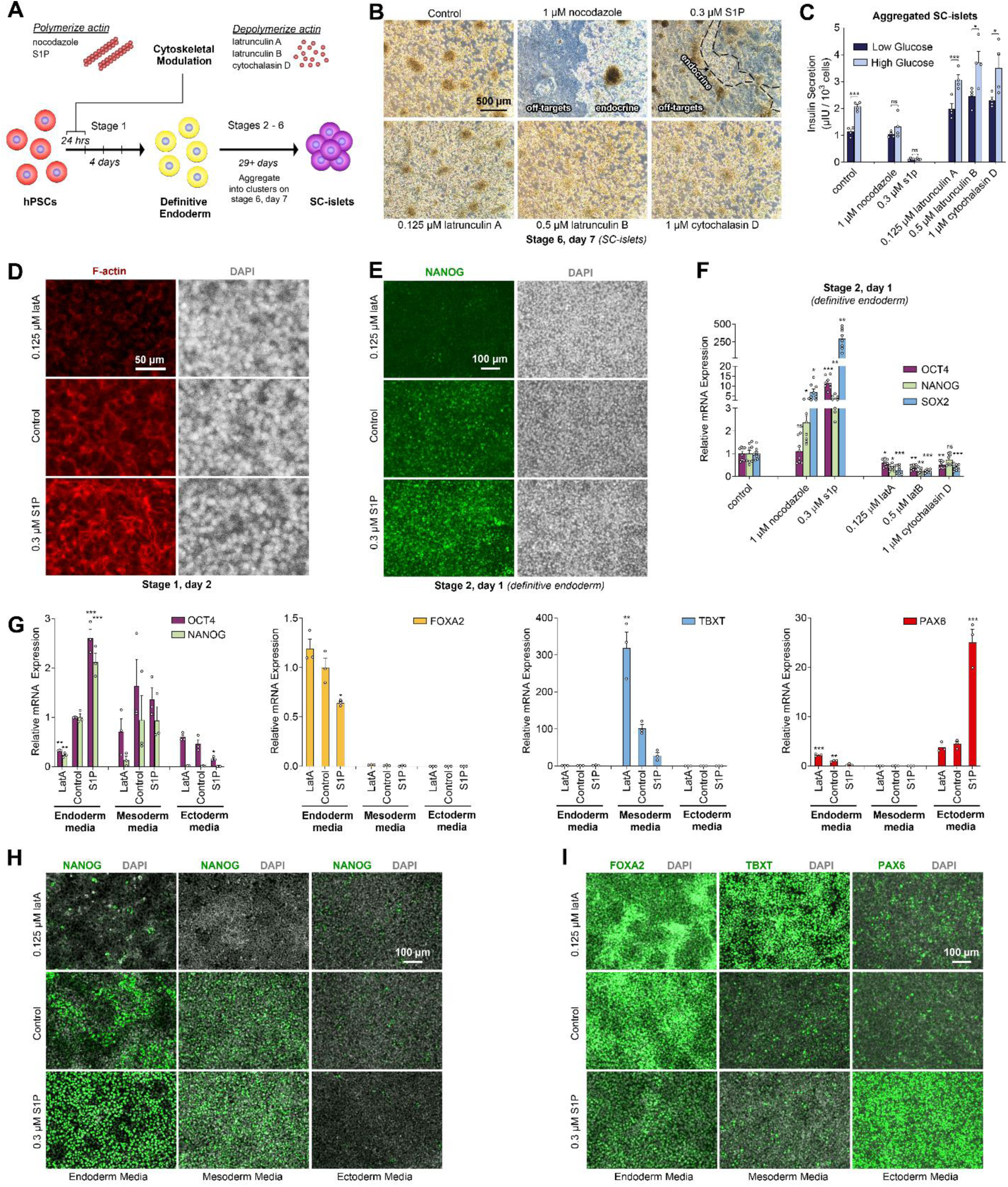
Actin polymerization state at the onset of differentiation influenced stem cell specification to all three germ layers. (**A**) Diagram of experiments testing cytoskeletal-modulating compounds during the SC-islet differentiation protocol. (**B**) Images of cells on stage 6, day 7 of this differentiation process, demonstrating that compounds which induce polymerization of the actin cytoskeleton (nocodazole, S1P) often produced non-endocrine cell types, while those that depolymerized the cytoskeleton (latrunculin A, latrunculin B, cytochalasin D) led to robust endocrine formation. Compounds were only added for the first 24 hours of differentiation. Scale bar = 500 µm. (**C**) Static GSIS of stage 6 cells treated with cytoskeletal-modulating compounds for the first 24 hours of differentiation (two-way paired t-test comparing insulin secretion between low and high glucose stimulation; n = 4). (**D**) Immunostaining images of F-actin on stage 1, day 2 in response to a 24-hour treatment with either latA, S1P, or no treatment. Scale bar = 50 µm. (**E**) Immunostaining images of NANOG at the end of stage 1 in response to either latA, S1P, or no treatment for the first 24 hours of differentiation. Scale bar = 100 µm. (**F**) qRT-PCR of pluripotency genes at the end of stage 1 in response to treatment with cytoskeletal-modulating compounds for the first 24 hours of differentiation (Welch’s ANOVA followed by Dunnett’s T3 multiple comparison test compares each condition to control; n = 8). (**G**) qRT-PCR of pluripotency genes and markers specific to each germ layer after differentiation in either endoderm, mesoderm, or ectoderm media. 0.125 µM latA or 0.3 µM S1P was added to the media during the first 24 hours of differentiation as indicated (ANOVA followed by Dunnett’s T3 multiple comparison test compares each condition to the control within each differentiation media; n = 3). Immunostaining of (**H**) NANOG and (**I**) lineage specific markers after this tri-lineage germ layer differentiation. Scale bars = 100 µm. All data were obtained with the HUES8 cell line. All data are represented as the mean, and all error bars represent the SEM. Individual data points are shown for all bar graphs. NS, not significant; *P *<* 0.05, **P *<* 0.01, ***P *<* 0.001.

In agreement with the improved SC-islet differentiation, cells receiving compounds that depolymerized F-actin (Fig. 1D, Supplemental Fig. 1A) displayed reduced expression of pluripotency markers (*OCT4*, *NANOG*, *SOX2*) by the end of stage 1 (Fig. 1E-F), indicating that these cells experienced a more robust exit from pluripotency. In contrast, compounds that induced actin polymerization (Fig. 1D, Supplemental Fig. 1A) exhibited much higher levels of these pluripotency genes (Fig. 1E-F), the extent of which was dependent upon the compound and degree of actin polymerization. While we initially tried to polymerize actin directly with jasplakinolide, we observed extensive cell death before reaching concentrations that visibly increased actin polymerization. Thus, we utilized compounds that induced actin polymerization indirectly through other mechanisms. Specifically, nocodazole depolymerizes microtubules, which has been shown to induce actin hypercontractility and polymerization.^44,45^ Sphingosine-1-phosphate (S1P), on the other hand, is a sphingolipid that has been shown to increase actin polymerization via activation of the Rho family of GTPases.^46,47^ Nocodazole specifically led to increased cortical F-actin as the cells rounded up after microtubule depolymerization, while S1P led to a more robust overall increase in actin polymerization (Fig. 1D, Supplemental Fig. 1A). Consequently, S1P-treated cells displayed the highest expression of pluripotency genes at the end of stage 1 (Fig. 1E-F), the most difficulty in forming FOXA2+/SOX17+ definitive endoderm (Supplementary Fig. 1B), and subsequently the fewest number of endocrine cells by the end of the protocol (Fig. 1B-C, Supplemental Fig. 1C-D).

Because the polymerization state of the actin cytoskeleton appeared to alter the exit of stem cells from pluripotency during directed differentiation to definitive endoderm, we tested its influence on the specification to the other germ layers using their respective differentiation protocols. Using a commercially available tri-lineage differentiation kit, we first replicated these results for the differentiation of stem cells to definitive endoderm. Specifically, depolymerizing F-actin with latA for the first 24 hours of differentiation decreased expression of pluripotency genes by the end of this definitive endoderm protocol, while increasing actin polymerization with S1P both maintained expression of pluripotency genes and reduced expression of the endoderm marker FOXA2 (Fig. 1G-I). We observed similar results with the directed differentiation to mesoderm, where latA treatment reduced pluripotency gene expression while simultaneously increased expression of the important mesoderm marker TBXT (Fig. 1G-I). In contrast, inducing actin polymerization with S1P during a directed ectoderm protocol greatly increased the expression of the key ectoderm marker PAX6 and reduced pluripotency gene expression (Fig. 1G-I). Thus, controlling the polymerization state of the actin cytoskeleton during the first 24 hours of differentiation influenced the exit of stem cells from pluripotency and could drastically improve key marker expression of each germ layer, but these effects were context dependent. Specifically, a depolymerized actin cytoskeleton at the onset of differentiation promoted mesendoderm lineages, while a polymerized actin cytoskeleton facilitated directed ectoderm differentiation.

### Depolymerizing the actin cytoskeleton during the first 24 hours of differentiation improved the specification to and function of SC-islets

To further explore the benefits of depolymerizing the actin cytoskeleton during the onset of directed differentiation to SC-islets, we tested a range of latA concentrations for the first 24 hours of stage 1 and continued the remainder of the SC-islet protocol as normal (Fig. 2A).^27^ It is important to note that this stage 1 latA treatment is in addition to the latA treatment at the beginning of stage 5 that we previously demonstrated to be crucial to the conversion of pancreatic progenitors into endocrine cells.^26^ The addition of the same compound at two separate time points during a directed stem cell differentiation protocol can lead to two different responses depending on cell state and signaling context. In this case, increasing latA concentration during the first 24 hours of stage 1 progressively induced a round morphology (Supplemental Fig. 2A) and disrupted actin polymerization during this initial 24-hour period (Supplemental Fig. 2B). Once latA was removed, the cells flattened out again for the rest of stage 1 (Supplemental Fig. 2C-D), as they were once again able to polymerize their actin cytoskeleton to pull against the tissue culture substrate. Interestingly, there was consistently more cell death throughout the rest of stage 1 even after latA was removed, particularly on days 3 and 4. However, a monolayer of cells remained attached to the plate throughout stage 1 at these latA concentrations (Supplemental Fig. 2C-D). If the latA concentration was too high, however, cell density ended up being too low by the end of stage 1, and these cells were unable to make endocrine cells by the end of the protocol (Supplemental Fig. 2E).

**Figure 2:**
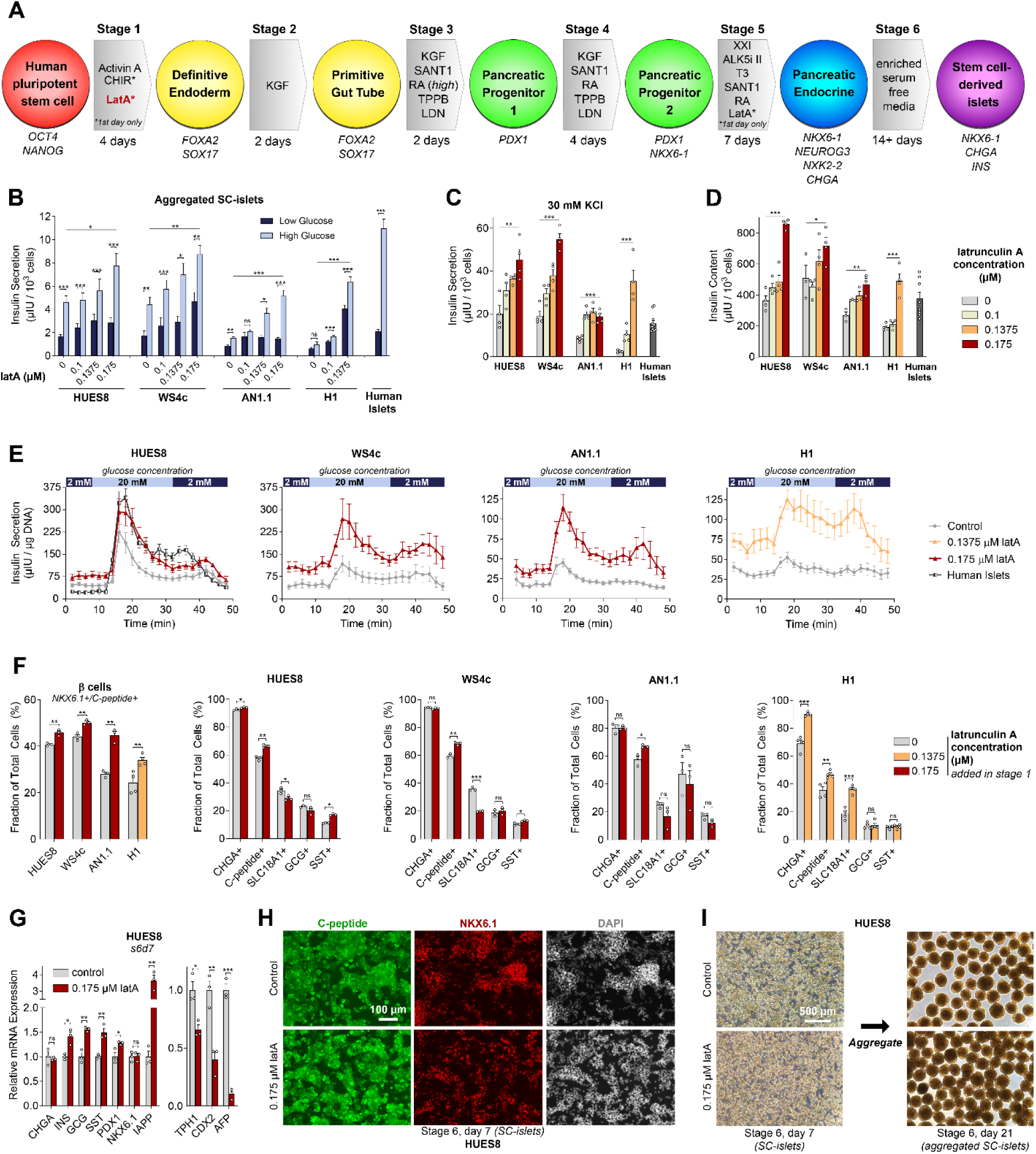
Latrunculin A treatment at the onset of definitive endoderm formation improves SC-islet function and composition by the end of the directed differentiation protocol. (**A**) Schematic of the SC-islet differentiation protocol used for all experiments incorporating the latA treatment during the first 24 hours of stage 1. (**B**) Static GSIS of stage 6 cells treated with latA (0 µM, 0.1 µM, 0.1375 µM, or 0.175 µM) during the first 24 hours of differentiation (two-way paired t-test comparing insulin secretion between low and high glucose stimulation; one-way ANOVA of high glucose between all latA concentrations for each cell line; n = 15-16 for HUES8, n = 6-8 for WS4c, n = 6-7 for AN1.1, n = 8 for H1, n = 11 for human islets from 2 separate donors). (**C**) Insulin secretion in response to 30 mM KCl stimulation in stage 6 cells treated with latA (0 µM, 0.1 µM, 0.1375 µM, or 0.175 µM) during the first 24 hours of differentiation (legend in (**D**); one-way ANOVA of insulin secretion between all latA concentrations for each cell line; n = 4 for all cell lines, n = 11 for human islets from 2 separate donors). (**D**) Insulin content of stage 6 cells treated with latA (0 µM, 0.1 µM, 0.1375 µM, or 0.175 µM) during the first 24 hours of differentiation (one-way ANOVA of insulin content between all latA concentrations for each cell line; n = 4 for HUES8, WS4c, and H1, n = 3 for AN1.1, n = 8 for human islets from 2 separate donors). (**E**) Dynamic insulin secretion of stage 6 cells treated with latA (0 µM, 0.1375 µM, or 0.175 µM) for the first 24 hours of differentiation (n = 7-10 for HUES8, n = 9 for WS4c, n = 6 for AN1.1, n = 6-8 for H1, n = 14 for human islets from 2 separate donors). (**F**) Flow cytometry on stage 6, day 7 demonstrated an increase in the percentage of SC-β cells (NKX6.1+/C-peptide+) for all cell lines in response to the stage 1 latA treatment and a decrease in the percentage enterochromaffin cells (SLC18A1+) for HUES8 and WS4c cell lines (two-way unpaired t-test, n = 3 for HUES8, WS4c, and AN1.1, n = 4 for H1). (G) qRT-PCR of stage 6, day 7 cells generated from HUES8 (two-way unpaired t-test, n = 3). (H) Immunostaining images of stage 6, day 7 cells generated from HUES8 stained for C-peptide and NKX6.1. Scale bar = 100 µm. (**I**) Images of stage 6, day 7 cells generated from HUES8 before and 2 weeks after aggregation into clusters. Scale bar = 500 µm. All data are represented as the mean, and all error bars represent the SEM. Individual data points are shown for all bar graphs. NS, not significant; *P *<* 0.05, **P *<* 0.01, ***P *<* 0.001.

We evaluated 4 hPSC lines with this latA treatment. Based on our prior work, the HUES8 and WS4c lines reliably generate high-functioning SC-islets, while differentiation from the AN1.1 and H1 lines is less efficient with our previous protocol.^27^ In this study, we found that 0.175 µM seemed to be the highest latA concentration that consistently resulted in high quality SC-islets by the end of the protocol for the HUES8, WS4c, and AN1.1 stem cell lines. The H1 stem cell line was more sensitive to this stage 1 latA treatment, and thus a maximum of 0.1375 µM latA was used for all H1 experiments. Strikingly, this latA treatment during the first 24 hours of the differentiation led to a dose-dependent increase in the function of SC-islets generated from these 4 hPSC lines by the end of the 5-week protocol. Specifically, this treatment drastically increased insulin secretion in both static (Fig. 2B) and dynamic (Fig. 2E) glucose-stimulated insulin secretion (GSIS) assays as well as in response to various secretagogues (Fig. 2C, Supplemental Fig. 3A). These SC-islets also exhibited much higher insulin content (Fig. 2D) while retaining a similar proinsulin to insulin content ratio (Supplemental Fig. 3B). With these improvements, the SC-islets approached the insulin secretion levels of primary human islets in response to glucose stimulation, particularly with the HUES8 and WS4c lines, and exceeded it in response to KCl stimulation for all 4 cell lines at the highest latA concentration. Similarly, SC-islets from all 4 cell lines had higher insulin content on a per cell basis than primary human islets at the highest latA concentration.

Upon further characterization, we observed that this stage 1 latA treatment consistently increased the number of SC-β cells by approximately 5-10% for any given batch (Fig. 2F), and this effect was dose-dependent (Supplemental Fig. 3C). For the HUES8, WS4c, and AN1.1 cell lines, the total number of CHGA+ endocrine cells did not change with the latA treatment, while the number of SCL18A1+ enterochromaffin cells decreased by approximately 5-15% (Fig. 2F). Thus, the stage 1 latA treatment improved the ratio of SC-β cells and the off-target enterochromaffin cells with these cell lines. Furthermore, the expression of genes associated with β cells (e.g, *INS*, *IAPP*) increased, while those associated with off-target lineages (e.g., *TPH1* for enterochromaffin cells*, CDX2* for intestinal cells*, AFP* for liver cells) decreased (Fig. 2G, Supplemental Fig. 3F,H,J). In the H1 line, the latA treatment appeared to also increase overall endocrine differentiation efficiency as indicated by an increase in the total number of CHGA+ cells, and thus there were also more SLC18A1+ cells in addition to the increase in SC-β cells (Fig. 2F). Re-normalizing the GSIS results to the number of SC-β cells rather than the total number of cells in the SC-islets (Supplemental Fig. 3D) revealed that latA-treated cells still exhibited higher insulin secretion, demonstrating that the stage 1 latA treatment not only increased the number of SC-β cells but also improved the insulin secretion per SC-β cell.

### Stage 1 latA treatment rescued differentiation to SC-islets from inconsistent hPSC lines

Throughout the protocol, the morphology of the latA-treated cells appeared more homogenous than the control (Supplemental Fig. 4A-B), though both conditions for all 4 hPSC lines could generate high quality endocrine cells that stained for SC-β cell markers (Fig. 2H, Supplemental Fig. 3E,G,I) and could be aggregated into islet-like clusters (Fig. 2I, Supplemental Fig. 5A-B). SC-islet yields with this stage 1 latA treatment were similar to our previously reported values for this differentiation process,^27^ typically generating between 0.5 and 0.75 million total cells per cm^2^ (Supplemental Figure 5C). The HUES8 and WS4c lines always yielded successful differentiations both with and without the stage 1 latA treatment, consistent with our previous publications demonstrating that these lines reliably generate high quality SC-islets that can cure severe pre-existing diabetes in mice without any signs of tumor formation. ^26,48,49^ Differentiations of the AN1.1 and H1 lines without the stage 1 latA treatment were inconsistent, however, often generating a mix of off-target and endocrine cell types or occasionally not producing any endocrine cells at all (Fig. 3A-B, Supplemental Fig. 6A-C). The H1 line has also been noted by others for being inefficient at differentiating to SC-islets.^50^ Strikingly, the stage 1 latA treatment rescued batches of cells in which the control differentiation failed to generate endocrine cells (Fig. 3A-C, Supplemental Figure 6A-C). This effect was dose-dependent, with the highest latA concentrations restoring the typical percentage (∼30-50%) of SC-β cells generated by the end of the protocol (Fig. 3C, Supplemental Fig. 6E). Interestingly, the control differentiations for the AN1.1 and H1 lines retained much higher expression of pluripotency genes by the end of stage 1 (Fig. 3D), mimicking the phenotype observed after treatment with actin-polymerizing compounds (Fig. 1F). These data suggest that the AN1.1 and H1 hPSC lines normally have trouble exiting pluripotency at the onset of differentiation, resulting in inefficiencies in pancreatic specification later on. In contrast, the latA treatment drastically decreased expression of *OCT4*, *NANOG*, and *SOX2* by the end of stage 1 in all four hPSC lines, further suggesting that the latA treatment helps hPSCs exit pluripotency more robustly (Fig. 3D).

**Figure 3:**
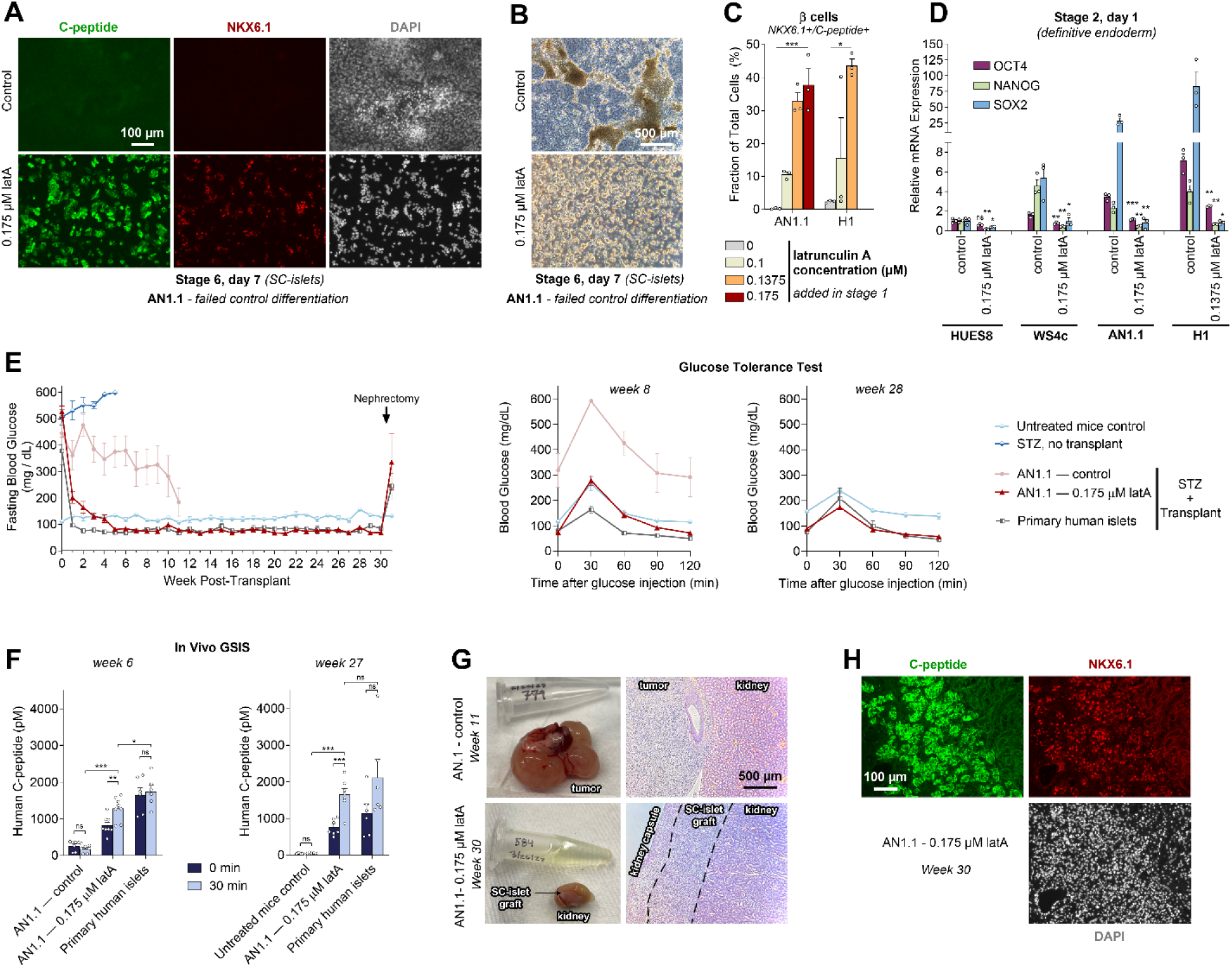
Latrunculin A treatment at the onset of definitive endoderm formation generates improved SC-islets from the AN1.1 cell line that can rapidly cure severe pre-existing diabetes in mice. (**A**) Immunostaining images of stage 6, day 7 cells generated from AN1.1 stained for C-peptide and NKX6.1. These images show a batch of cells that failed to make endocrine cells with the normal differentiation protocol, while treating this same batch of cells with 0.175 µM latA during stage 1 rescued SC-β cell generation. Scale bar = 100 µm. (**B**) Images on stage 6, day 7 showing a batch of AN1.1 cells that failed to make endocrine cells with the normal differentiation protocol but exhibited the expected endocrine morphology when treated with 0.175 µM latA during stage 1. Scale bar = 500 µm. (**C**) Flow cytometry on stage 6, day 7 of differentiation batches from the AN1.1 and H1 cell lines in which the control differentiation failed to produce SC-β cells, while SC-β cell generation was rescued by treating these same batches with latA during stage 1 in a dose-dependent manner (one-way ANOVA, n = 3). (**D**) qRT-PCR of pluripotency genes on stage 2, day 1 cells for all four cell lines, with and without stage 1 latA treatment (two-sided t-tests between control and latA condition for each gene in a given cell line, n = 3). (**E**) Mice were made diabetic with streptozotocin (STZ) injections. Cells were transplanted underneath the kidney capsule using either primary human islets or SC-islets generated with the AN1.1 stem cell line. These SC-islets were produced either without or with the 0.175 µM latA treatment during stage 1 (untreated control: n = 5 mice; STZ, no transplant: n = 7 mice; AN1.1 – control: n = 7 mice; AN1.1 – 0.175 µM latA: n = 8 mice; primary human islets: n = 6 mice). Fasting blood glucose levels demonstrated rapid restoration of blood glucose control with the mice transplanted with the primary human islets and the AN1.1 – 0.175 µM latA SC-islets. These transplanted mice also controlled blood glucose levels during glucose tolerance tests at 8 and 28 weeks. (**F**) In vivo GSIS assays at weeks 6 and 27 demonstrated that the transplanted AN1.1 – 0.175 µM latA SC-islets secreted more insulin 30 minutes after glucose injection, and this insulin secretion was at comparable levels to the transplanted human islets by week 27 (two-way paired t-test between low and high glucose stimulation; two-way unpaired t-test of the 30 minute timepoint between the mice transplanted with AN1.1 – 0.175 µM latA SC-islets and either the AN1.1 – control SC-islets or human islets). (**G**) The mice transplanted with the AN1.1 – control cells formed large tumors by week 11, while the mice transplanted with the AN1.1 – 0.175 µM latA SC-islets did not have any signs of overgrowths during the experiment. Furthermore, the AN1.1 – 0.175 µM latA SC-islet graft was still clearly visible in the kidney after 30 weeks. (**H**) Many SC-β cells (NKX6.1+/C-peptide+) were observed in these AN1.1 –0.175 µM latA SC-islet grafts after 30 weeks in the mice. All data are represented as the mean, and all error bars represent the SEM. Individual data points are shown for all bar graphs. NS, not significant; *P *<* 0.05, **P *<* 0.01, ***P *<* 0.001.

Off-target or residual undifferentiated cells produced during SC-islet differentiations as a result of inefficient exit from the pluripotent state are an important safety concern for a cell-based therapy, as they could form tumors upon transplantation.^51,52^ To further demonstrate the drastic improvement that this stage 1 latA treatment could have on SC-islet generation, function, and safety, we transplanted SC-islets generated both with and without the stage 1 latA treatment using the AN1.1 hPSC line under the kidney capsule of severely diabetic mice. SC-islets generated with this induced pluripotent stem cell (iPSC) line have not been previously transplanted into mice. The fasting blood glucose levels of the mice receiving the SC-islets generated without the latA treatment (AN1.1 – control) came down gradually, finally dipping below 200 mg/dL by week 11 (Fig. 3E). In contrast, the fasting blood glucose of the mice transplanted with the latA-treated SC-islets (AN1.1 – 0.175 µM latA) fell below 200 mg/dL between weeks 1 and 2 (Fig. 3E). By week 5, the fasting blood glucose of these mice was approximately 80 mg/dL, matching the blood glucose levels of mice transplanted with primary human islets (Fig. 3E). The fasting blood glucose of the mice transplanted with the AN1.1 –0.175 µM latA SC-islets remained between approximately 70-80 mg/dL for the remainder of the experiment (30 weeks total), mimicking the fasting blood glucose set point in humans. Similarly, random blood glucose measurements of these mice averaged 146 mg/dL by week 6 and came down further overtime (Supplemental Fig. 7A). A glucose tolerance test at week 8 demonstrated that the AN1.1 – 0.175 µM latA SC-islets were much better at controlling blood glucose levels than the control cells (Fig. 3E). Furthermore, they secreted much more insulin and were glucose responsive *in vivo* (Fig. 3F), mirroring the large difference observed during *in vitro* GSIS (Fig. 2B). As they matured overtime *in vivo*, the AN1.1 – 0.175 µM latA SC-islets secreted more insulin (Fig. 3F), became more glucose responsive (Fig. 3F), and improved their ability to control blood glucose to a level similar as primary human islets (Fig. 3E). The kidneys containing the AN1.1 – 0.175 µM latA SC-islets and primary human islets were removed 30 weeks after transplantation, resulting in a rapid reversal to a diabetic state in these mice (Fig. 3E). These grafts looked healthy with no overgrowths and contained many C-peptide+/NKX6.1+ SC-β cells (Fig. 3G-H), similar to the primary human islet grafts (Supplemental Fig. 7B). In contrast, the AN1.1 – control SC-islets formed tumors by week 11 (Fig. 3G), despite transplanting batches of cells that successfully generated SC-islets. This tumor formation is unsurprising, however, given the observed off-targets that were often generated with the AN1.1 line without latA treatment (Supplemental Fig. 6C). Taken together, these data demonstrate that treating the otherwise inconsistent AN1.1 hPSC line with latA during the first 24 hours of stage 1 generated SC-islets with drastically improved function that were able to rapidly cure severely diabetic mice to a similar degree as in our previous publications using the HUES8 and WS4c lines^26,48,49^ and without tumor formation.

### Stage 1 latA treatment altered the signaling dynamics of major pathways during the specification of definitive endoderm

Collectively, these data suggest that cytoskeletal state is critically important for the exit of hPSCs from pluripotency and during definitive endoderm specification, ultimately influencing the subsequent identity and function of differentiated SC-islets. To further elucidate the mechanism by which latA influenced this process, we performed single cell RNA-sequencing every day of stage 1 on HUES8 cells treated with or without 0.175 µM latA for the first 24 hours of differentiation (Fig. 4A). After these initial 24 hours, the latA-treated cells had decreased expression of multiple genes encoding metallothionein proteins (*MT1H*, *MT1G*, *MT2A*, *MT1E*, *MT1F*) as well as increased expression of several genes related to histone structure (*HIST1H1B*, *HIST1H1D*), inhibition of canonical WNT signaling (*DKK1*, *DKK4*), and stress mediation (*TXNIP*) (Supplemental Fig. 8A). Expression of actin (*ACTB*) itself also decreased in response to the latA treatment (Supplemental Fig. 8A-B). Despite differential expression of these few genes, the control and latA-treated cells clustered similarly for the first several days of stage 1 (Fig. 4A). As stage 1 progressed, however, the latA-treated cells became more distinct, separating from the untreated cells after stage 1, day 3 (s1d3). By the end of stage 1, the latA-treated cells formed their own distinct cluster with notable differences in the expression of genes known to be involved in definitive endoderm formation, while the control cells at s2d1 clustered more similarly to the previous day (Fig. 4A-B).

**Figure 4:**
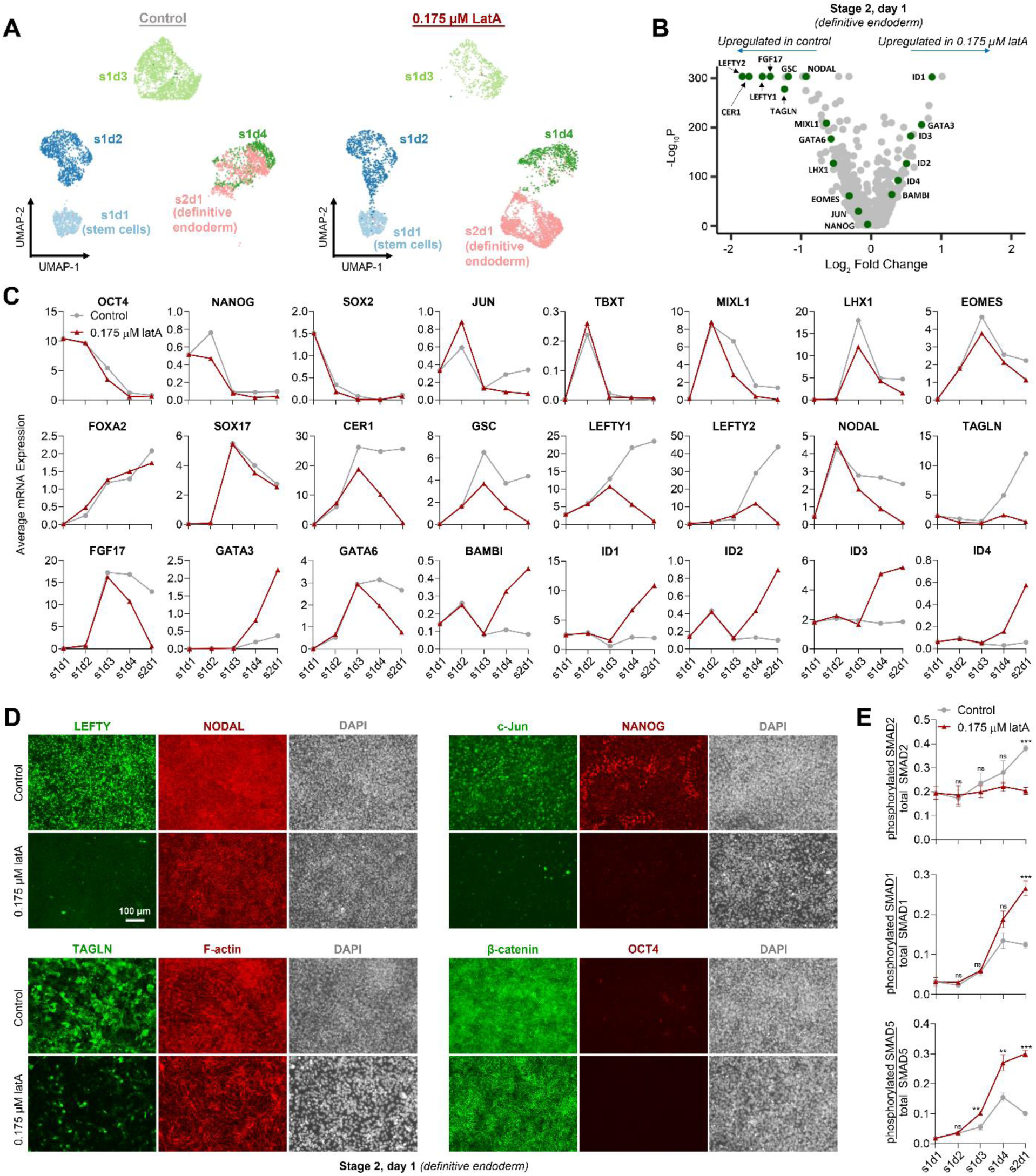
Latrunculin A treatment alters the signaling dynamics of key pathways during endoderm formation. (**A**) Single cell RNA-sequencing UMAPs of cells every day of stage 1 comparing without and with a 0.175 µM latA treatment for the first 24 hours of differentiation. Example nomenclature: s1d1 = stage 1, day 1. (**B**) Volcano plot of genes differentially expressed at the end of stage 1 with latA treatment. (**C**) The average gene expression of all cells in a cluster from the single cell RNA-sequencing data at each day of differentiation either without or with latA treatment. (**D**) Immunostaining images of several important markers at the end of stage 1, confirming the expression patterns exhibited at the transcriptional level in the single cell RNA-sequencing data. Scale bar = 100 µm. (**E**) Enzyme-linked immunosorbent assays (ELISAs) measuring the ratio of phosphorylated to total protein of SMADS 1, 2, and 5 every day of stage 1 without or with latA treatment (two-way unpaired t-test at each time point, n = 4). All assays were performed with the HUES8 cell line. All data are represented as the mean, and all error bars represent the SEM. NS, not significant; *P *<* 0.05, **P *<* 0.01, ***P *<* 0.001.

We compared the gene expression of the control and latA-treated cells throughout stage 1. The average expression of the pluripotency markers *OCT4*, *NANOG*, and *SOX2* decreased as expected, though they appeared to turn off more rapidly in latA-treated cells (Fig. 4C, Supplemental Fig. 8C). The mesendoderm markers *TBXT* and *MIXL1* peaked on s1d2 but were downregulated on subsequent days as the cells differentiated into *FOXA2*+/*SOX17*+ definitive endoderm (Fig. 4C, Supplemental Fig. 8C). While the expression of *FOXA2* and *SOX17* progressed similarly between the control and latA-treated cells throughout stage 1, other markers of definitive endoderm exhibited unique expression patterns. Specifically, while *CER1* and *GSC* expression peaked on s1d3 in both conditions, their expression was strongly downregulated by the end of stage 1 in the latA-treated cells but maintained in the control cells (Fig. 4C, Supplemental Fig. 8C). Similarly, *LHX1*, *EOMES*, and *JUN* were more downregulated by the end of stage 1 in the latA-treated cells (Fig. 4C, Supplemental Fig. 8C).

Identifying some of the most differentially expressed genes at s2d1 revealed further interesting trends of genes related to several major signaling pathways known to be involved in the specification of definitive endoderm (Fig. 4B). Specifically, while the expression levels of the TGF-β superfamily members *NODAL* and its antagonists *LEFTY1* and *LEFTY2* initially rose in both conditions, they were strongly downregulated in the latA-treated cells by the end of stage 1 (Fig. 4C, Supplemental Fig. 8C). *TAGLN*, which encodes the actin-binding protein transgelin and is downstream of TGF-β signaling, concurrently rose in the control cells (Fig. 4C, Supplemental Fig. 8C). Conversely, the expression of the TGF-β inhibitor *BAMBI* and the BMP downstream targets *ID1*, *ID2*, *ID3*, and *ID4* all increased drastically during the second half of stage 1 (Fig. 4C, Supplemental Fig. 8C). The expression of several other genes further distinguished the latA-treated population by the end of stage 1, including increased *GATA3* expression and downregulated *GATA6* and *FGF17* expression (Fig. 4C, Supplemental Fig. 8C). Re-clustering of the cells only at s2d1 highlighted the distinct population generated by the latA-treated cells by the end of stage 1 when compared to the control cells (Supplemental Fig. 9A). Despite most cells expressing the definitive endoderm markers *FOXA2* and *SOX17* in both conditions, the expression of many of these key genes were constrained to either population (Supplemental Fig. 9B). qRT-PCR revealed similar expression patterns for these genes in all 4 hPSC lines in response to the stage 1 latA treatment (Supplementary Fig. 8D). In a parallel experiment, treating hPSCs with latA in mTeSR1 altered expression of many of these same genes (e.g., *NODAL*, *LEFTY1*, *LEFTY2*, *JUN*, *TAGLN*, *ID1*, and multiple metallothionein genes), highlighting the strong effect latA has on the expression of these specific genes (Supplemental Fig. 10A-B). Interestingly, however, a much higher latA concentration was required in mTeSR1 to see the same rounding effect on cell morphology when compared to latA added in stage 1 media, both with and without Activin A and CHIR99021 (Supplemental Fig. 10C).

In agreement with the gene expression data, single-nuclei ATAC sequencing at the end of stage 1 demonstrated that the control cells had higher promoter chromatin accessibility for *NODAL* (Supplementary Fig. 11B,D). Furthermore, the control cells had higher motif chromatin accessibility for the SMAD2::SMAD3::SMAD4 and FOS::JUN binding domains, indicative of increased signaling through the Activin/Nodaland JNK-JUN pathways, respectively (Supplemental Fig. 11B-C). Conversely, latA-treated cells had greater motif accessibility for *HNF1A* and *HNF1B* (Supplemental Fig. 11B-C), which are important during the specification of the gut tube that occurs after definitive endoderm formation. Taken together, these data indicate that while the latA treated cells generated *SOX17*+/ *FOXA2*+ definitive endoderm in a similar proportion as the untreated control cells, they formed a unique endodermal state that was better at generating functional SC-β cells (Fig. 2). This unique endodermal state appears to be the result of drastic changes in the temporal dynamics of several key signaling pathways in the latA-treated cells, including increased BMP signaling and decreased Activin/Nodal signaling during the second half of stage 1.

To confirm these signaling changes at the protein level, we immunostained for a number of these markers throughout stage 1. Mirroring the gene expression patterns, LEFTY1/2 and NODAL initially increased in both conditions but were downregulated during the second half of stage 1 in latA-treated cells (Fig. 4D, Supplemental Fig. 12A-C). C-Jun stained much more intensely on s1d2 in the latA-treated cells but was strongly downregulated by the end of stage 1 in these cells, while the control cells retained c-Jun protein throughout stage 1 (Fig. 4D, Supplemental Fig. 13B-C). Furthermore, latA-treated cells had essentially eliminated NANOG and OCT4 protein by the end of stage 1, while heterogeneous NANOG protein expression and a few OCT4 positive cells persisted in the control (Fig. 4D, Supplemental Fig. 13A,C). Similar to the trends observed in gene expression, the amount of transgelin protein increased as stage 1 progressed in the control but not in latA-treated cells (Fig. 4D, Supplemental Fig. 14B-C). While latA-treated cells recovered their ability to polymerize actin after latA removal, differences in F-actin staining persisted throughout stage 1 (Fig. 4D, Supplemental Fig. 13A,C). Mimicking the pattern of F-actin staining, β-catenin became much more localized to the cell membrane in latA-treated cells, suggesting a difference in canonical WNT signaling (Fig. 4D, Supplemental Fig. 15A-B). Another interesting observation from these immunostaining studies was that, despite all hPSC lines initially staining homogenously for OCT4 and NANOG to demonstrate pluripotency (Supplemental Fig. 16A), there was heterogeneity in initial c-Jun and F-actin staining between hPSC lines. Specifically, H1 and AN1.1 hPSCs appeared to have lower overall c-Jun staining than either HUES8 or WS4c, and AN1.1 hPSCs in particular had heterogenous F-actin staining (Supplemental Fig. 16B).

As further evidence for these changes in stage 1 signaling dynamics, the downstream Activin/Nodal signaling effector SMAD2 had a decreased percentage of proteins in their phosphorylated state with the latA treatment during the second half of stage 1, indicating decreased Activin/Nodal signaling during this time (Fig. 4E). Conversely, the BMP signaling effectors SMAD1 and SMAD5 had an increased percentage of phosphorylated proteins during this time period in latA-treated cells, suggesting increased BMP signaling in the second half of stage 1 (Fig. 4E). In support of the notion that BMP signaling is important to definitive endoderm formation and downstream specification of the pancreas, application of the BMP inhibitor LDN193189 from s1d2 through the end of stage 1 prevented the formation of PDX1+/NKX6.1+ pancreatic progenitors by s5d1, even in latA-treated cells (Supplemental Fig. 17A-B). The dynamics of WNT signaling throughout stage 1 also seemed to be different in latA-treated cells. For example, while the ratio of phosphorylated GSK-3β to the total amount of GSK-3β protein was lower in latA-treated cells for most of stage 1, it increased sharply during the last day, indicating changes in canonical WNT signaling during this time (Supplemental Fig. 17D). Total β-catenin protein followed a similar trend (Supplemental Fig. 17D), and gene expression of WNT signaling components changed during this time as well (Supplemental Fig. 17C). Collectively, these data demonstrate that treating cells with latA during the first 24 hours of differentiation influenced the signaling dynamics throughout stage 1 of major pathways known to be involved in the specification of definitive endoderm, generating a unique definitive endoderm population by the end of stage 1.

### Stage 1 latA treatment altered downstream patterning of the gut tube and improved pancreatic progenitor specification

Despite these differences in major signaling pathways during stage 1, both conditions yielded close to 100% FOXA2+/SOX17+ definitive endoderm by the end of stage 1 (Fig. 5A). By s5d1, however, the latA-treated cells had a higher percentage of PDX1+/NKX6.1+ cells (Fig. 5B-C), indicating better specification to pancreatic progenitors. Furthermore, the latA-treated cells demonstrated higher expression of pancreatic progenitor genes (*PDX1*, *NKX6.1*, *PTF1α, SOX9*) as well as decreased expression of genes associated with other endodermal lineages (*CDX2*, *OTX2, AFP, HNF4a*) (Fig. 5D). To further elucidate the changes in signaling induced by the latA treatment that led to improved pancreatic progenitor specification and subsequent SC-β cell formation, we performed single cell RNA-sequencing at each stage of differentiation (Fig. 5E). Overall, the control and latA-treated cells clustered similarly at most stages, with the end of stage 1 (s2d1, definitive endoderm) still exhibiting the most distinct populations between the two conditions. Despite these similarities, the two conditions still demonstrated differences in the expression of important genes at various stages (Fig. 5F). Specifically, markers of anterior foregut (*OTX2*, *LHX1, FZD8, SISHA2*, *SIX3*, *HESX1*) were elevated in the control cells at s2d1 and s3d1 (Fig. 5F, Supplemental Fig. 18B). By s4d1, however, the control cells demonstrated elevated levels of posterior markers (*AFP*, *TTR*), while the latA-treated cells exhibited transient upregulation of more anterior markers (*SOX2, OSR1*) (Fig. 5F, Supplemental Fig. 18B). By s5d1, importantly, the latA-treated cells demonstrated higher expression of key pancreatic markers (*ONECUT1*, *PTF1a*, *PDX1*, *NKX6.1*) (Fig. 5F, Supplemental Fig. 18B). Confirming some of these changes at the protein level, we observed that the amount of the anterior marker OTX2 was initially higher at s2d1 in the control (Fig. 5H, Supplemental Figs. 19A-B). Conversely, the latA-treated cells had higher protein levels of PDX1 and ONECUT1 by s4d1 (Fig. 5H, Supplemental Figs. 19A-C) as well as NKX6.1 by s5d1 (Fig. 5C). Re-clustering of the sequencing data at s6d7 demonstrated that the latA-treated cells generated an increased percentage of β cells and a lower percentage of enterochromaffin cells in the final populations (Fig. 5J), further confirming that these improvements in pancreatic progenitor specification resulted in improved β cell differentiation. Furthermore, the β cell population demonstrated higher expression of β cell genes (e.g., *INS*, *IAPP*, *ERO1B*) and lower expression of other hormones (*GCG*) as well as genes not normally found in β cells (*LDHA*, *DDC*) (Supplemental Fig. 18D).

**Figure 5:**
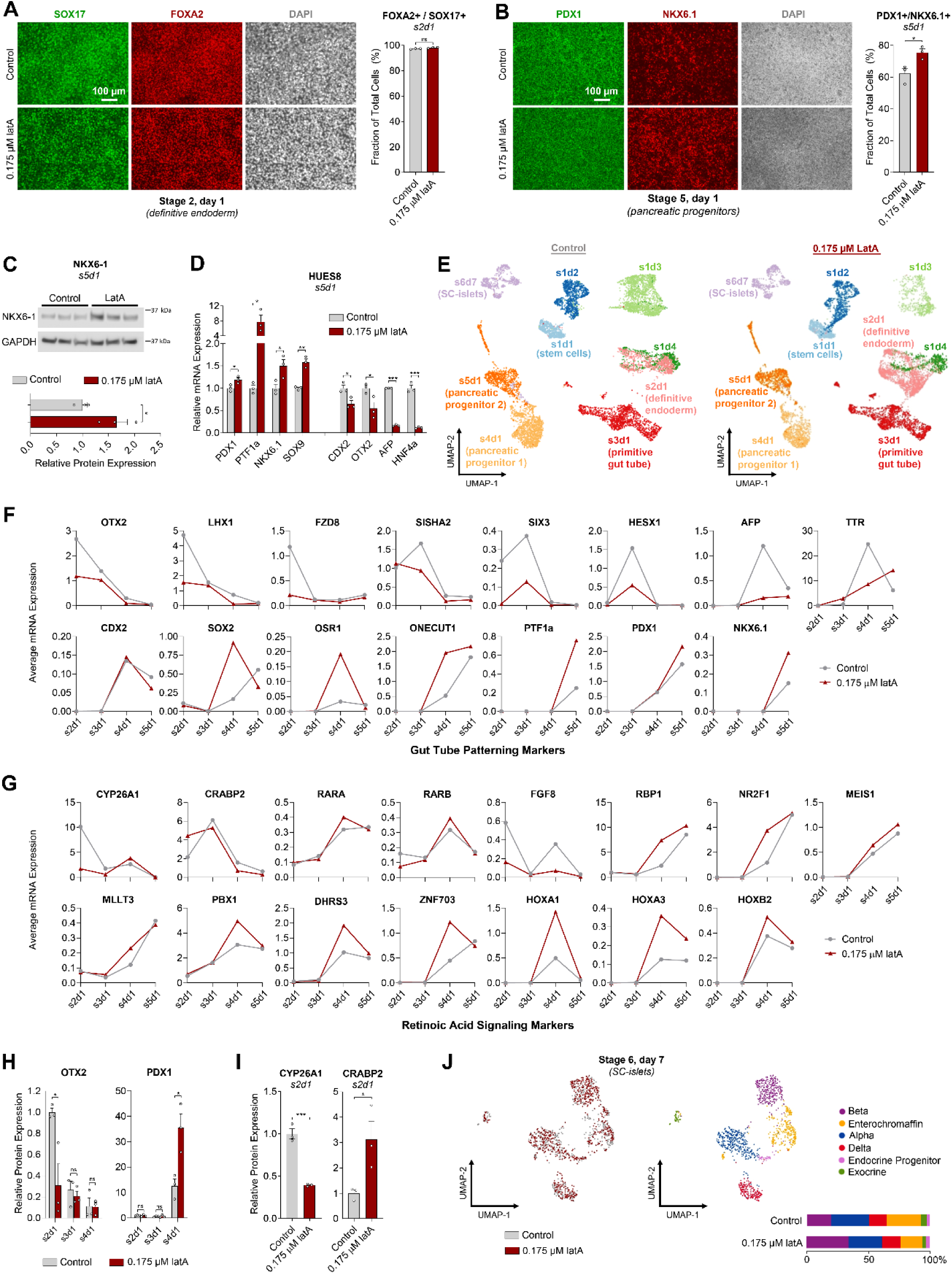
Latrunculin A treatment at the onset of definitive endoderm formation alters the expression of lineage-specific genes as well as those associated with RA signaling during subsequent gut tube patterning. (**A**) Immunostaining images and flow cytometry (two-way unpaired t-test, n = 3) of the definitive endoderm markers FOXA2 and SOX17 at the end of stage 1 comparing without and with a 0.175 µM latA treatment for the first 24 hours of differentiation. Scale bar = 100 µM. (**B**) Immunostaining images and flow cytometry (two-way unpaired t-test, n = 3) of the pancreatic markers PDX1 and NKX6.1 at the end of stage 4 comparing without and with a 0.175 µM latA treatment for the first 24 hours of differentiation. Scale bar = 100 µM. (**C**) Quantified western blots of the NKX6.1 protein at the end of stage 4 comparing without and with a 0.175 µM latA treatment for the first 24 hours of differentiation (two-way unpaired t-test, n = 3). (**D**) qRT-PCR at the end of stage 4 for pancreatic markers (*PDX1*, *PTF1α*, *NKX6.1*, *SOX9*) and markers associate other lineage of the gut tube (*CDX2*, *OTX2*, *AFP*, *HNF4a*) (two-way unpaired t-test, n = 3). (**E**) Single cell RNA-sequencing UMAPs of cells throughout the SC-islet differentiation process comparing without and with a 0.175 µM latA treatment for the first 24 hours of differentiation. Example nomenclature: s1d1 = stage 1, day 1. (**F-G**) The average gene expression of all cells in a cluster from the single cell RNA-sequencing data at the intermediate stages of differentiation (s2d1, s3d1, s4d1, and s5d1) either without or with latA treatment. (**H-I**) Quantification of western blots for the indicated proteins comparing without and with a 0.175 µM latA treatment for the first 24 hours of differentiation (two-way unpaired t-test, n = 3). (**J**) Single cell RNA-sequencing UMAPs of cells on stage 6, day 7 of differentiation, demonstrating differences in the percentage of cell types in latA-treated cells. All assays were performed with the HUES8 cell line. All data are represented as the mean, and all error bars represent the SEM. Individual data points are shown for all bar graphs. NS, not significant; *P *<* 0.05, **P *<* 0.01, ***P *<* 0.001.

Other genes that stood out in this single cell RNA-sequencing dataset for exhibiting differences in the latA-treated cells during differentiation were those relating to components of the retinoic acid (RA) signaling pathway as well as its downstream targets. In particular, *CYP26A1*, which degrades RA, was strongly elevated in the control cells at s2d1 (Fig. 5G, Supplemental Fig. 18C). In contrast, *CRABP2*, which helps shuttle RA to the nucleus and promote RA signaling, was elevated at this time in latA-treated cells (Fig. 5G, Supplemental Fig. 18C). These differences were confirmed at the protein level (Fig. 5I, Supplemental Fig. 17B) and suggest that latA-treated cells may be better primed for RA signaling in subsequent stages. During stage 3, a high dose of RA is used to induce specification of the posterior portion of the foregut and expression of the key pancreatic transcription factor PDX1 (Supplemental Fig. 18A). Expectedly, the nuclear RA receptors RARA and RARB peaked at the end of this stage (s4d1) (Fig. 5G, Supplemental Fig. 18C). Interestingly, the latA-treated cells exhibited higher expression of many RA target genes at s4d1 (*RBP1*, *NR2F1*, *MEIS1*, *MLLT3*, *PBX1*, *DHRS3*, *ZNF703*, *HOXA1*, *HOXB2*, *HOXA3*),^53–56^ indicating increased RA signaling during stage 3 (Fig. 5G, Supplemental Fig. 18C). These included several of the HOX genes, which are important for segmenting the gut tube into different organ-forming regions.^34,37^ Correspondingly, the latA-treated cells had increased PDX1 protein expression at s4d1 (Fig. 5H, Supplemental Fig. 19A-B), indicating better specification of the pancreatic portion of the gut tube. These peaks in expression of RA-associated genes also coincided with the peaks of other important gut tube genes mentioned earlier (Fig. 5F-G, Supplemental Fig. 18B-C). Conversely, the control cells demonstrated increased expression of *FGF8* at this timepoint (Fig. 5G, Supplemental Fig. 18C). Because RA signaling is known to repress *FGF8* expression,^56,57^ this further suggests that the control cells experienced decreased RA signaling during stage 3. Taken together, these data indicate the latA treatment in stage 1 influenced the temporal expression of key genes in subsequent stages that altered gut tube patterning, resulting in better specification of pancreatic progenitors by s5d1 and improved SC-β cell differentiation by the end of the protocol. These effects appear to be at least partially mediated by altered RA signaling dynamics during the specification of the posterior foregut.

To further characterize SC-islets generated with the stage 1 latA treatment, we performed single-nucleus multi-omic sequencing (mRNA and ATAC) on s6d15 after the cells had been aggregated into islet-like clusters for over one week (Fig. 6A, Supplemental Fig. 20A-C). Similar to the observations made with the single cell RNA-sequencing and flow cytometry analysis on s6d7, the combined RNA and ATAC characterization demonstrated that the SC-islet clusters generated with the stage 1 latA treatment had a higher percentage of β cells and a lower percentage of enterochromaffin cells (Fig. 6A). Furthermore, each cell type in the SC-islets generated with the stage 1 latA treatment generally demonstrated a more mature identity for that cell type in terms of both mRNA expression (Fig. 6B) as well as promoter accessibility for those genes (Fig. 6C). Looking specifically at the β cell population, the latA-treated cells had higher mRNA expression of genes such as *INS* and those involved in calcium regulation (*CACNA1A, ASPH*) (Fig. 6D). They also exhibited increased promoter accessibility for β cell genes (*INS*, *ISL1, ONECUT1*) as well as multiple HOX genes (Fig. 6D). Furthermore, the CTCF binding motif was more accessible in the control, which we previously demonstrated to promote an enterochromaffin lineage (Fig. 6D).^58^ In summary, treating the cells with latA for the first 24 hours of differentiation altered the signaling dynamics during definitive endoderm formation, leading to altered gut tube patterning and better pancreatic progenitor specification. These cells ultimately generated SC-islets that had more β cells as well as a decreased number of enterochromaffin cells. Importantly, these β cells exhibited a more mature identity and secreted more insulin in response to glucose stimulation.

**Figure 6.**
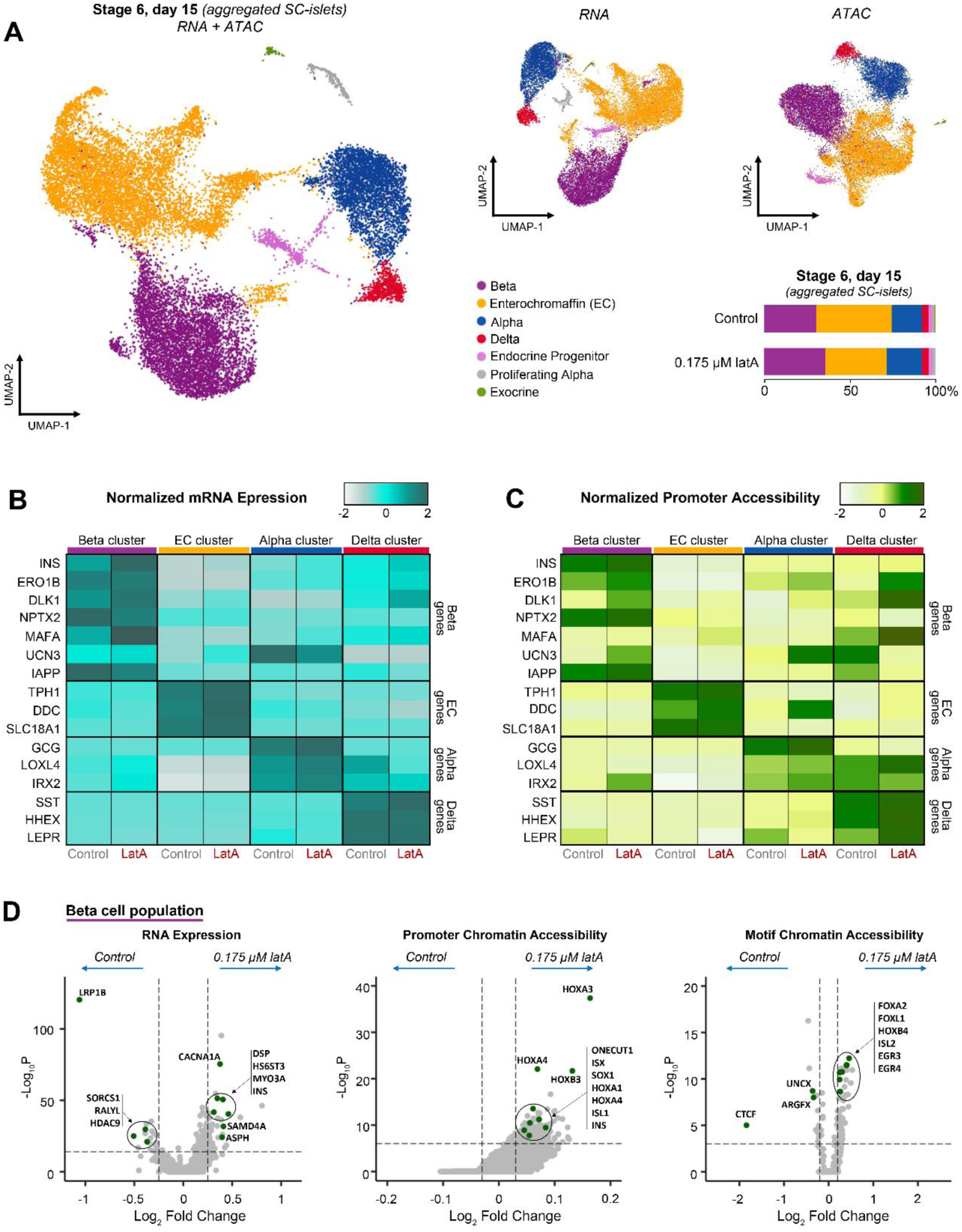
Latrunculin A treatment during the first 24 hours of differentiation generates SC-islets containing cells with more mature cell identities. (**A**) Single-nuclei multi-omic sequencing UMAPs showing cell clustering based on either gene expression, chromatin accessibility, or both combined from each individual cell. Cells were processed on stage 6, day 15, which was 8 days after aggregation into islet-like clusters. Treating cells with 0.175 µM latA during the first 24 hours of stage 1 influenced the percentages of the different cell types produced by the end of the protocol. (**B**) Gene expression heatmap of markers associated with different endocrine cell types observed in the multi-omic sequencing. (**C**) Heatmap of the chromatin accessibility for the promoters of these gene markers. (**D**) Volcano plots within the β cell cluster showing differences in gene expression, chromatin accessibility for different gene promoters, and the chromatin accessibility of transcription factor binding motifs in SC-islets generated without and with 0.175 µM latA for the first 24 hours of differentiation. All multi-omic sequencing was performed with the HUES8 cell line.

## DISCUSSION

In this study, we demonstrated that the polymerization state of the actin cytoskeleton at the onset of differentiation influenced the exit of stem cells from pluripotency and could drastically improve key marker expression during differentiation to each of the three germ layers in a context-dependent manner. In particular, a depolymerized cytoskeleton at the onset of differentiation promoted mesendoderm lineages in their respective differentiation media, while a polymerized actin cytoskeleton facilitated directed ectoderm differentiation. Applying these findings specifically to an SC-islet differentiation protocol, we demonstrated that depolymerizing the actin cytoskeleton with latA during the first 24 hours of differentiation facilitated a more robust exit from pluripotency and altered signaling dynamics during definitive endoderm formation, modifying the temporal activation of the Activin/Nodal, BMP, JNK-JUN, and WNT pathways. These changes in stage 1 signaling influenced downstream patterning of the gut tube, including modified expression of components of the RA signaling pathway and altered HOX gene expression. Subsequently, latA treated cells exhibited improved pancreatic identity by s4d1 and decreased expression of genes associated with other endodermal lineages along the gut tube, such as *AFP* (liver) and *CDX2* (intestine). This improved pancreatic specification led to an increased percentage of PDX1+/NKX6.1+ pancreatic progenitors by the end of stage 4 and ultimately generated SC-islets with a higher percentage of β cells and lower percentage of enterochromaffin cells by the end of the protocol. These SC-islets contained β cells with a more mature identity and greatly improved insulin secretion in response to glucose stimulation. Furthermore, this latA treatment could rescue differentiations from inconsistent hPSC lines in batches that otherwise failed to produce endocrine cells.

While the exact mechanistic link between actin polymerization state and these specific changes in signaling dynamics remains to be fully elucidated, the data suggest that depolymerizing the actin cytoskeleton with latA helped transition the hPSCs out of pluripotency during differentiation to definitive endoderm. For example, compounds that increased actin polymerization caused these cells to retain higher expression of pluripotency genes by the end of stage 1 (Fig. 1F), and consequently these cells had difficulty differentiating into SC-islets (Fig. 1B-C). Similarly, the AN1.1 and H1 hPSC lines retained much higher expression of these pluripotency genes at the end of stage 1 in the absence of the latA treatment, which correlated with the inconsistent differentiations normally observed with these hPSC lines. Even with the HUES8 hPSC line, the control cells demonstrated heterogenous NANOG protein expression and some OCT4 positive cells by the end of stage 1, while these proteins were essentially gone in the latA-treated cells (Fig. 4D, Supplemental Fig. 13A,C). Additionally, the HUES8 control cells consistently appeared to lag behind the latA-treated cells throughout the differentiation. For instance, the anterior marker OTX2 was turned off more slowly in the control during stages 2 and 3, while the master pancreatic regulator PDX1 turned on more strongly in the latA-treated cells in stage 3 (Fig. 5H, Supplemental Fig. 19A-B).

These results support emerging evidence for the importance of cytoskeletal dynamics in pluripotent stem cells.^42^ Interestingly, some of the only genes strongly downregulated immediately after the 24-hour latA treatment during our SC-islet protocol were those encoding metallothionein proteins, which play an important role in the homeostasis of metal ions within a cell (Supplemental Fig. 8A, 10A). In agreement with this observation, it has been demonstrated that metallothionein genes must be downregulated in order for cells to differentiate into definitive endoderm.^59^ Furthermore, differences in metallothionein expression may contribute to variation in pluripotency between stem cell lines.^60^ Similarly, JNK-JUN signaling has also been shown inhibit exit from pluripotency and impede differentiation to definitive endoderm.^61^ While latA-treated cells exhibited a strong initial increase in nuclear accumulation of c-Jun, c-Jun was quickly eliminated in the subsequent days from latA-treated cells as they regained their ability to polymerize actin (Fig. 4C-D, Supplemental Fig. 13B-C). In agreement with this observation, depolymerizing actin has been shown to strongly increase c-Jun expression.^62^ Thus, depolymerization of F-actin at the onset of differentiation appears to affect the timing and intensity of JNK-JUN signaling, which may subsequently influence how cells exit pluripotency and respond to differentiation factors. Interestingly, we observed differences in initial c-Jun staining in the H1 and AN1.1 hPSC lines as well as a heterogenous F-actin structure in AN1.1 (Supplemental Fig. 16B). These observations suggest that differences in endogenous signaling involving actin and c-Jun may contribute to the difficulties these hPSC lines had in exiting pluripotency and subsequently differentiating into SC-islets without latA treatment.

Because F-actin depolymerization influenced hPSC exit from pluripotency during our SC-islet differentiation protocol, the signaling dynamics changed during stage 1, leading to a unique definitive endoderm population by the end of this stage. Many of these signaling differences centered around the Activin/Nodal and BMP signaling pathways, which are critical during early embryogenesis.^31–33^ Our results indicate that a definitive endoderm population with these unique gene signatures is able to more robustly make functional SC-islets. In particular, the latA-treated cells appeared to be more receptive to RA signaling during the subsequent patterning of the gut tube, leading to better specification of pancreatic progenitors.

The improvements in SC-islet generation described here have noteworthy implications for their performance and safety as a cell therapy for treating diabetes. Prior studies have demonstrated that SC-islets produced from hPSCs are immature *in vitro* in terms of both function and identity.^22,23,28,43,50,58,63,64^ Here, we were able to significantly improve insulin secretion, insulin content, and cell identity of SC-islets with multiple cell lines. This new process increased the proportion of SC-β cells and lowered the percentage of enterochromaffin cells in SC-islets, demonstrating improved pancreatic specification and reduction of unwanted off-target cell types. Furthermore, this stage 1 latA treatment increased differentiation consistency and reduced line-to-line variability, particularly with the H1 and AN1.1 hPSC lines. For example, while we have never observed tumor formation when transplanting SC-islets generated with the HUES8 and WS4c lines with our recent protocols,^26,48,49^ residual off-target cells in the control AN1.1 differentiations proliferated and caused tumors in mice by week 11 after transplantation (Fig. 3G). In contrast, the stage 1 latA treatment of AN1.1 cells generated SC-islets that both rapidly cured severely diabetic mice and prevented tumor formation (Fig. 3E-H). This reversal of hyperglycemia was maintained long-term in mice without evidence of hypoglycemia. Thus, this new version of the SC-islet differentiation protocol can help improve the performance of SC-islets while simultaneously preventing the formation of troublesome off-target cell types, particularly when utilizing new cell lines. The ability to seamlessly adapt SC-islet differentiation protocols to clinical grade cell lines is crucial for the safety, efficacy, and speed of translation of this cell-based therapy. Furthermore, this new differentiation strategy could be combined with other advances in transplantation to develop novel cell therapy products.^65–68^

In terms of the practicality of using this new method, latA can simply be added for the first 24 hours of our previously published SC-islet differentiation protocol.^27^ The latA concentrations for the cell lines shown here can be used as written. However, when starting with a new cell line, we recommend testing a range of concentrations, with the 0.1 µM to 0.175 µM range being a good place to start. The concentration of latA needed is dependent upon the particular cell line and starting cell density. The goal should be to get the cells as round as possible after 24 hours without too many of them falling off the plate (Fig. 1D, Supplemental Fig. 2A-B), as the protocol will not work if cell density becomes too low (Supplemental Fig. 2E). If the cells are very round on the second day of differentiation, the media can be changed several hours early to prevent too many cells from detaching from the plate. Of particular note, there will be more cell death throughout stage 1 than without the latA treatment, especially on s1d3 and s1d4 when the media can be cloudy with floating cells. As long as there is a confluent layer of cells attached to the flask, however, the differentiation should be successful (Supplemental Fig. 2C-D). While this latA treatment improved differentiation to SC-islets with the 4 hPSC lines presented here, it is possible that it will not work for all hPSC lines, particularly if those lines have variations in endogenous signaling of some of the pathways influenced by the latA treatment. For cell lines that are very sensitive to latA, latrunculin B or cytochalasin D can be tried as alternatives, as cells treated with these compounds also generated SC-islets with improved function.

It is important to note that this stage 1 latA treatment is in addition to the latA treatment on s5d1, and the alteration of cytoskeletal state seems to be influencing different pathways at their respective points in the differentiation. In directed stem cell differentiation protocols, timing is extremely important. Adding the same compound at two separate time points can lead to two different responses depending on cell state and signaling context, as is the case here. Interestingly, the latA concentration used during stage 1 had to be much lower than the concentration used later on in the protocol during stage 5. Furthermore, a much higher concentration of latA was needed in mTeSR1 than in stage 1 media in order to induce the same rounding effect on hPSC morphology (Supplemental Fig. 10C), emphasizing the context-dependent effect of latA on cytoskeletal signaling.

In summary, our results highlight the critical importance of cytoskeletal state at the onset of directed stem cell differentiation protocols. Specifically, we demonstrated that polymerizing actin during the first 24 hours of directed differentiation promoted ectoderm specification, whereas depolymerizing F-actin during this initial 24-hour window improved specification to mesendoderm lineages. We successfully applied this concept to a pancreatic differentiation protocol to generate SC-islets with greatly improved identity and function. Thus, we anticipate that carefully optimizing the polymerization state of the actin cytoskeleton at the onset of differentiation could improve the specification of other cell types during their respective directed differentiation protocols.

## METHODS

Detailed, step-by-step instructions for most of the methods described here can be found in our previous publication.^27^

### Stem cell culture

Two human embryonic stem cell (hESC) lines (HUES8 [RRID: CVCL_B207] and H1 [RRID: CVCL_9771]) and two induced pluripotent stem cell (iPSC) lines (WS4c [RRID: CVCL_A9K6] and AN1.1 [RRID: CVCL_A9K3]) were used in this study, as we have described previously.^27^ The hESC lines were used under the approval of the Washington University Embryonic Stem Cell Research Oversight Committee (approval no. 15-002) with the appropriate consent and conditions. To propagate the undifferentiated stem cells, they were seeded onto Matrigel (356230, Corning) coated tissue culture-treated flasks at 100,000 cells/cm^2^ in mTeSR1 (85850, Stemcell Technologies) supplemented with 10 µM Y-27632 (ab120129, Abcam) and cultured in a humidified incubator at 5% CO_2_ and 37°C. The media was replaced every day with new mTeSR1, increasing the volume added each day as the cells grew. After 3-4 days, the cells were passaged using 0.2 mL TrypLE Express/cm^2^ (12604-039, Gibco) and seeded again in mTeSR1 supplemented with 10 µM Y-27632 for either further propagation or for differentiation.

### SC-islet differentiation

The full detailed protocol used to differentiate the stem cells into SC-islets as well as assays to assess their quality can be found in our previous publication.^27^ In brief, stem cells were seeded onto Matrigel-coated tissue culture-treated 6-well plates or flasks (T75 or T182.5) in mTeSR1 supplemented with 10 µM Y-27632 at the following densities: HUES8 = 0.63 cells/cm^2^, WS4c and H1 = 0.53 cells/cm^2^, AN1.1 = 0.74 cells/cm^2^. After 24 hours, differentiation was initiated by replacing the mTeSR1 with Stage 1 media and factors. The cells were fed every day with the following media:

#### Base media formulations

<u>BE1</u>: 500 ml MCDB 131 (10372019, Gibco) supplemented with 0.8 g glucose (G7528, MilliporeSigma), 0.587 g sodium bicarbonate (S5761, MilliporeSigma), 0.5 g BSA (68700, Proliant Biologicals) and 5 mL GlutaMAX (35050-079, Gibco). <u>BE2</u>: 500 mL MCDB 131 supplemented with 0.4 g glucose, 0.587 g sodium bicarbonate, 0.5 g BSA, 5 mL GlutaMAX and 22 mg vitamin C (A4544, MilliporeSigma). <u>BE3</u>: 500 mL MCDB 131 supplemented with 0.22 g glucose, 0.877 g sodium bicarbonate, 10 g BSA, 2.5 mL ITS-X (51500-056, Gibco), 5 ml GlutaMAX and 22 mg vitamin C. <u>S5</u>: 500 mL MCDB 131 supplemented with 1.8 g glucose, 0.877 g sodium bicarbonate, 10 g BSA, 2.5 ml ITS-X, 5 mL GlutaMAX, 22 mg vitamin C, 5 mL penicillin/streptomycin solution (30-002-CI, Corning) and 5 mg heparin (H3149, MilliporeSigma). <u>Enriched Serum Free Media (ESFM)</u>: 500 mL MCDB 131 supplemented with 0.23 g glucose, 10.5 g BSA, 5.2 ml GlutaMAX, 5.2 mL penicillin/streptomycin solution, 5 mg heparin, 5.2 mL MEM nonessential amino acids (20-025-CI, Corning), 84 μg ZnSO4 (10883, MilliporeSigma), 523 μL Trace Elements A (25-021-CI, Corning) and 523 μL Trace Elements B (25-022-CI, Corning).

#### Differentiation media

<u>Stage 1 (4 days)</u>: BE1 medium supplemented with 100 ng/mL Activin A (338-AC, R&D Systems) and 3 μM CHIR99021 (2520691, Peprotech) for the first 24 hours, followed by 3 days of BE1 containing 100 ng/mL Activin A only. <u>Stage 2 (2 days)</u>: BE2 medium supplemented with 50 ng/mL KGF (AF-100-19, Peprotech). <u>Stage 3 (2 days)</u>: BE3 medium supplemented with 50 ng/ml KGF, 200 nM LDN193189 (SML0559, MilliporeSigma), 500 nM TPPB (53431, Tocris Bioscience), 2 μM retinoic acid (R2625, MilliporeSigma) and 0.25 μM SANT1 (S4572, MilliporeSigma). <u>Stage 4 (4 days)</u>: BE3 medium supplemented with 50 ng/mL KGF, 200 nM LDN193189, 500 nM TPPB, 0.1 μM retinoic acid and 0.25 μM SANT1. <u>Stage 5 (7 days)</u>: S5 medium supplemented with 10 μM ALK5i II (ALX-270-445-M005, Enzo Life Sciences), 1 μM T3 (64245, MilliporeSigma), 1 μM XXI (565790, MilliporeSigma), 0.1 μM retinoic acid, and 0.25 μM SANT1. 1 µM Latrunculin A (10010630, Cayman Chemical) was added to this medium for the first 24 hours only. <u>Stage 6 (7–25 days)</u>: Cultures were kept on the plate with ESFM for up to 7 days. To generate islet-like clusters, cells were single-cell dispersed with TrypLE and resuspended in 6 mL of ESFM within a 6-well plate at a concentration of 5-8 million cells per well on an orbital shaker (Orbi-Shaker CO_2_, Benchmark Scientific) at 115 RPM. Functional assessments were performed 7-18 days after cluster aggregation.

#### Differentiation nomenclature

Throughout the manuscript, the differentiations are labeled with both the stage and the day number in that stage to indicate the specific point of the process that data was collected. Furthermore, data was always taken before new media were added that day. For example, cells at stage 1, day 1 (s1d1) were stem cells that were plated for 24 hours but that had not yet received the first day of differentiation media. Similarly, cells at stage 2, day 1 (s2d1) were cells still in stage 1 differentiation media immediately before changing to stage 2 differentiation media.

#### Cytoskeletal compound screen

To investigate the influence of cytoskeletal state on SC-islet differentiation, the following compounds were added to stage 1 media at the same time as AA and CHIR99021 on s1d1: 0.25 µM jasplakinolide (11705, Cayman Chemical), 1 µM nocodazole (13857, Cayman Chemical), 0.3 µM S1P (62570, Cayman Chemical), 0.125 µM latA (10010630, Cayman Chemical), 0.5 µM latrunculin B (10010631, Cayman Chemical), and 1 µM cytochalasin D (C2618, MilliporeSigma). On the following day, this media was replaced with BE1 supplemented with 100 ng/mL Activin A only. The rest of the differentiation was continued as previously described.^27^

#### Differentiations with latA

For differentiations testing the influence of latA added during stage 1, the specified latA concentration (either 0 µM, 0.1 µM, 0.1375 µM, or 0.175 µM) was added to stage 1 media at the same time as AA and CHIR99021 on s1d1. On the following day, this media was replaced with BE1 supplemented with 100 ng/mL Activin A only. The rest of the differentiation was continued as previously described^27^, with the following slight modifications. All cells were fed everyday of stage 5 rather than skipping days during the second half of this stage. Once the media was changed to ESFM at the onset of stage 6, the cells from a few cell lines tended to bud off the plate as clusters, particularly on days 3 and 4 of stage 6. If this occurred, the clusters were collected from the media and placed in 6-well plates on an orbital shaker. Furthermore, the aggregation step could be performed early, starting on day 4 of stage 6. Finally, the aggregated clusters generated with the stage 1 latA treatment had a tendency to stick together during culture. Thus, the orbital shaker speed was increased from 100 RPM to 115 RPM, which completely resolved this issue.

#### Differentiations with stage 1 BMP inhibition

For experiments testing BMP inhibition during stage 1, 200 nM LDN193189 was added to the stage 1 differentiation media starting on s1d2 through the end of stage 1. The remainder of the differentiation was continued normally.

### Tri-lineage differentiation

To differentiate hPSCs into the three different germ layers, HUES8 stem cells were plated onto Matrigel-coated 6-well plates in mTeSR1 supplemented with 10 µM Y-27632 at a density of 0.63 cells/cm^2^. After 24 hours, the media was replaced with either the endoderm, mesoderm, or ectoderm differentiation media from a commercially available kit (05230, Stemcell Technologies). Cells were fed according to the manufacturer’s instructions, with the exception that either 0.125 µM latA or 0.3 µM S1P was added to the media during the first 24 hours only. At the end of this differentiation stage, cells were either collected for RNA extraction or fixed for immunohistochemistry.

### Microscopy and immunocytochemistry

A Leica DMi1 inverted light microscope was used to take all brightfield images, while a Leica DMI4000 B inverted microscope captured all fluorescence images. To stain cells for fluorescent imaging, they were fixed with 4% paraformaldehyde (PFA) (157-4-100, Electron Microscopy Sciences) at room temperature for 30 minutes. The cells were then blocked and permeabilized for 45 minutes at room temperature with an immunocytochemistry (ICC) solution consisting of 0.1% Triton X (327371000, Acros Organics) and 5% donkey serum (017000-121, Jackson ImmunoResearch) in PBS (21-040-CV, Corning). These cells were then incubated with primary antibodies diluted in ICC solution overnight at 4 °C. After 24 hours, the cells were washed with ICC solution, incubated with secondary antibodies diluted in ICC solution for 2 hours at room temperature, and finally stained with DAPI nuclear stain (D1306, Invitrogen) for 15 minutes at room temperature.

For histological sectioning, excised mouse kidneys containing transplanted cells were fixed overnight with 4% PFA at 4°C. These samples were submerged in 70% ethanol and sent off to be paraffin embedded and sectioned by Histowiz, Inc (Long Island City, New York). Paraffin was removed from sectioned samples with Histo-Clear (C78-2-G, Thermo Fisher Scientific), and antigen retrieval was carried out in a pressure cooker (2100 Retriever, Electron Microscopy Sciences) with 0.1 M EDTA (AM9261, Ambion). Slides were blocked and permeabilized with ICC solution for 45 min and then incubated with primary antibodies in ICC solution overnight at 4 °C. After 24 hours, the slides were incubated with secondary antibodies for 2 hours at room temperature and then finally sealed with DAPI Fluoromount-G (0100-20, SouthernBiotech). For hematoxylin and eosin staining, sections were processed and stained by the Pulmonary Morphology Core at Washington University.

#### Antibodies for immunocytochemistry

Primary antibodies were diluted in ICC solution at 1:300 unless otherwise indicated: rat anti-C-peptide (GN-ID4-S, Developmental Studies Hybridoma Bank (DSHB)), 1:100 mouse anti-NKX6-1 (F55A12-S, DSHB), goat anti-PDX1 (AF2419, R&D Systems), mouse anti-SOX17 (MAB1924, R&D Systems), rabbit anti-FOXA2 (07-633, MilliporeSigma), mouse anti-OCT-3/4 (sc-5279, Santa Cruz Biotechnology), goat anti-NANOG (AF1997, R&D Systems), rabbit anti-Lefty (ab22569, Abcam), mouse anti-Nodal (ab55676, Abcam), rabbit anti-c-Jun (ab40766, Abcam), rabbit anti-TAGLN (ab14106, Abcam), rabbit anti-beta catenin (ab32572, Abcam), mouse anti-OTX2 (NBP2-37597, Novus Biologicals), sheep anti-ONECUT1 (AF6277, Novus Biologicals), rabbit anti-BRACHYURY (D2Z3J, Cell Signaling Technology), and rabbit anti-PAX6 (D3A9V, Cell Signaling Technology). F-actin was stained directly with 1:500 TRITC-conjugated phalloidin (FAK 100, MilliporeSigma).

Secondary antibodies were diluted in ICC solution at 1:300. All secondary antibodies were raised in donkey: anti-rat Alexa Fluor 488 (A21208, Invitrogen), anti-mouse Alexa Fluor 594 (A21203, Invitrogen), anti-rabbit Alexa Fluor 488 (A21206, Invitrogen), anti-goat Alexa Fluor 594 (A11058, Invitrogen), anti-mouse Alexa Fluor 488 (A21202, Invitrogen), anti-rabbit Alexa Fluor 594 (A21207, Invitrogen), anti-goat Alexa Fluor 488 (A11055, Invitrogen), and anti-sheep Alexa Fluor 594 (A11016, Invitrogen).

### Flow cytometry

Samples were single-cell dispersed with TrypLE for 10 minutes at 37°C, mixed with an equal volume of PBS, and centrifuged for 3 minutes at 300g. After removing the supernatant, 4% PFA was added to the tubes, which were immediately shaken vigorously to disperse the cell pellet. After fixing for 30 minutes at 4°C, the cells were washed once with PBS and incubated in ICC solution 45 minutes at 4°C. The cells were incubated with primary antibodies diluted in ICC solution overnight at 4°C, washed once with ICC solution, and then incubated in secondary antibodies diluted in ICC solution for 2 hours at 4°C. The cells were washed once with PBS and then run on a Cytek Northern Lights flow cytometer (Cytek Biosciences). Results were analyzed with FlowJo (version 10.8.1).

#### Antibodies for flow cytometry

Primary antibodies were added to ICC solution at the indicated dilutions: 1:300 rat anti-C-peptide (GN-ID4-S, Developmental Studies Hybridoma Bank (DSHB)), 1:100 mouse anti-NKX6-1 (F55A12-S, DSHB), 1:300 goat anti-PDX1 (AF2419, R&D Systems), 1:1,000 rabbit anti-CHGA (ab15160, Abcam), 1:250 mouse anti-SST conjugated to Alexa Fluor 488 (566032, BD Biosciences), 1:350 mouse anti-GCG conjugated to BV421 (565891, BD Biosciences), 1:300 rabbit anti-SCL18A1 (HPA063797, MilliporeSigma), 1:625 mouse anti-FOXA2 conjugated to PE (561589, BD Biosciences), 1:625 anti-SOX17 conjugated to APC (130-111-033, Miltenyi Biotech).

Secondary antibodies were diluted in ICC solution at 1:500 for flow cytometry. All secondary antibodies were raised in donkey: anti-rat PE (712-116-153, Jackson ImmunoResearch), anti-mouse Alexa Fluor 488 (A21202, Invitrogen), anti-rabbit Alexa Fluor 647 (A31573, Invitrogen), anti-goat Alexa Fluor 647 (A21447, Invitrogen), and anti-mouse Alexa Fluor 647 (A31571, Invitrogen).

### qRT-PCR

mRNA was extracted using the RNeasy Mini Kit (74016, Qiagen) according to the manufacturer’s instructions, and samples were treated with DNAse (79254, Qiagen) during extraction. cDNA was synthesized with the High Capacity cDNA Reverse Transcriptase Kit (4368814, Applied Biosystems). Quantitative real time PCR (qRT-PCR) was run on a QuantStudio 6 Pro (A43180, Applied Biosystems) and analyzed using the ΔΔCt method. *TBP* was used as the housekeeping gene. Primers can be found in Supplementary Table 1.

### GSIS

#### Static GSIS

Culture inserts (PIXP01250, MilliporeSigma) were placed in a 24-well plate and wetted with 500 µL of Krebs buffer (KrB) (128 mM NaCl, 5 mM KCl, 2.7 mM CaCl_2_, 1.2 mM MgSO_4_, 1 mM Na_2_HPO_4_, 1.2 mM KH_2_PO_4_, 5 mM NaHCO_3_ 10 mM HEPES (15630-080, Gibco) and 0.1% BSA). About 50 stage 6 SC-islet clusters were pipetted into each insert and washed twice with 1 mL KrB containing 2 mM glucose. The inserts were then transferred to new wells with 1 mL KrB containing 2 mM glucose and incubated at 37° C and 5% CO_2_ for 1 hour to equilibrate the samples at low glucose. To transfer each insert between wells, they were angled at approximately 45° until the liquid drained through the membrane from the insert into the well. After 1 hour, the supernatants were discarded, and the inserts were transferred into new wells with 1 mL KrB containing 2 mM glucose. After incubating for 1 hour, the inserts were transferred to new wells with 1 mL KrB containing 20 mM glucose. The supernatants were saved, as this was the insulin secreted at 2 mM glucose over the 1-hour incubation. After an additional hour, the inserts were transferred to new wells containing 1 mL of TrypLE to single-cell disperse the clusters and perform cell counts (Vi-Cell XR, Beckman Coulter). The supernatants were again saved, as this was the insulin secreted at 20 mM glucose over the 1-hour incubation. The supernatants at low and high glucose were quantified with a human insulin ELISA kit (80-INSHU-E10.1, ALPCO), and results were normalized to the cell count for each sample.

To measure insulin secretion in response to different secretagogues, one of the following compounds was added during the high glucose challenge: 30 mM KCl (BP366500, Thermo Fisher Scientific), 300µM tolbutamide (T0891, MilliporeSigma), or 100µM IBMX (I5879, MilliporeSigma).

#### Dynamic GSIS

Approximately 50 stage 6 SC-islet clusters were loaded into cell chambers (PERI-CHAMBER, BioRep Diabetes) by sandwiching them between two layers of hydrated Bio-Gel P-4 polyacrylamide beads (150-4124, Bio-rad). These chambers were attached to a high-precision eight-channel dispenser pump (ISM931C, Ismatec) via 0.015-inch inner diameter inlet and outlet tubing and 0.04-inch inner diameter connection tubing (PERI5-TUBSET-PVC, BioRep Diabetes). Dispensing nozzles were attached to the end of the outlet tubing (PERI-NOZZLE, BioRep Diabetes). The chambers were immersed in a 37°C water bath for the remainder of the assay. The samples were first equilibrated for 90 minutes with a KrB solution containing 2mM glucose at a flow rate of 100 µL per minute. After this first 90 minutes, the outflow was collected in 2-minute intervals for 48 minutes. The perfusing KrB solution contained 2 mM glucose for the first 8 minutes, 20 mM glucose for the next 24 minutes, and then back to 2 mM glucose for 16 minutes. The SC-islets were then lysed with a solution of 10 mM Tris (T6066, MilliporeSigma), 1 mM EDTA (AM9261, Ambion) and 0.2% Triton X (327371000, Acros Organics). DNA was quantified using the Quant-iT Picogreen dsDNA assay kit (P7589, Invitrogen) and was used to normalize insulin values quantified with a human insulin ELISA kit.

### Insulin and proinsulin content

Approximately 50 stage 6 clusters were pipetted into Eppendorf tubes and washed with PBS. In parallel, an equal amount of clusters were single-cell dispersed with TrypLE and counted. 500 µL of an acid-ethanol solution (1.5% HCl and 70% ethanol) were added to each tube, vortexed vigorously, and incubated at 20°C overnight. After 24 hours, the tubes were vortexed vigorously and incubated again at 20°C. After an additional 24 hours, the samples were centrifuged at 2100g for 15 minutes. The supernatants were collected and added to an equal volume of 1M TRIS solution (T2319, MilliporeSigma). These samples were quantified with a proinsulin ELISA kit (10-1118-01, Mercodia) and a human insulin ELISA kit. Proinsulin and insulin secretion values were normalized to the viable cell counts.

### Protein quantification

To isolate protein, cells were rinsed with ice-cold PBS and scraped from 6-well plates into Eppendorf tubes. The PBS was removed, and 0.5 mL of ice-cold cell lysis buffer (9803, Cell Signaling) was added. For the SMAD, GSK-3β, and total β-catenin ELISA assays, this lysis buffer was supplemented 10 µL/mL of a protease and phosphatase inhibitor cocktail (78440, Thermo Scientific). For western blots, 1mM phenylmethylsulfonyl fluoride (P-470-10, GoldBio) was added to the lysis buffer. The samples were incubated for 5 minutes on ice, sonicated for 30 seconds, and centrifuged for 10 minutes at 14,000g and 4°C. The supernatant was collected, and protein concentration was determined by BCA assay (23225, Pierce).

#### Western blots

20 µg of protein with 4x LDS buffer (2463559, Invitrogen) was boiled for 5 minutes at 95°C and separated on a 4-12% Bis-Tris gel (NP0321BOX, Invitrogen) in MES running buffer (NP0002, Invitrogen). The proteins were transferred onto PVDF membrane (1620177, BioRad), blocked in 5% milk (1706404, BioRad) for 1 hour at room temperature, and incubated in primary antibodies in 5% milk overnight at 4°C. The following day, blots were washed and incubated in secondary antibody for 1 hour at room temperature. Blots were run with protein standards (1610377, BioRad), exposed with chemiluminescence (1705060, Biorad), and imaged on a Licor Odyssey FC. All blots were normalized to GAPDH.

#### Antibodies for western blots

Primary antibodies were used at the indicated dilutions: 1:50 mouse anti-NKX6-1 (F55A12-S, DSHB), 1:300 goat anti-PDX1 (AF2419, R&D Systems), 1:500 mouse anti-OTX2 (NBP2-37597, Novus Biologicals), 1:300 mouse anti-CYP26A1 (AB151968, Abcam), 1:2000 rabbit anti-CRABP2 (ab211927, Abcam), and 1:2000 GAPDH (sc-32233, Santa Cruz Biotechnology).

#### SMAD ELISAs

ELISA kits measuring the ratio of phosphorylated to total SMAD1, 2, and 5 were carried out according to the manufacturer’s instructions (ab279928, ab279930, and ab279934, Abcam). 8 µg of protein was loaded for the SMAD 1 and 5 assays, while 16 µg protein was loaded for SMAD2 detection. Incubations and washes were followed as recommended, and absorbance was read at 450 nm (Biotek Synergy H1, Agilent).

#### *GSK-3*β *ELISAs*

ELISA kits measuring the amount of phosphorylated (7265C, Cell Signaling Technology) and total GSK-3β (7311C, Cell Signaling Technology) were carried out according to the manufacturer’s instructions. Samples were diluted to a concentration of 0.25 mg/mL, and 100 µL was loaded into each well of the assay plate. Incubations and washes were followed as recommended, and absorbance was read at 450 nm (Biotek Synergy H1, Agilent).

#### *Total* β-*catenin ELISA*

An ELISA kit was used to quantify the total amount of β-catenin according to the manufacturer’s instructions (ab275100, Abcam). Samples were diluted to a concentration of 300 µg total protein/mL, and 15 µg was loaded into each well of the assay plate. Incubations and washes were followed as recommended, and absorbance was read at 450 nm (Biotek Synergy H1, Agilent).

### Transplantation studies

All animal studies were performed in accordance with Washington University International Care and Use Committee (IACUC) regulations (protocol #21-0240). Prior to transplantation, 7 week-old male immunodeficient mice (NOD.Cg-Prkdc^scid^ Il2rg^tm1Wjl^/SzJ, Jackson Laboratories) were treated with 45 mg/kg streptozotocin (1621500, R&D Systems) in 0.9% sterile saline (51-405022.052, Moltox) for 5 consecutive days. Mice were diabetic (blood glucose > 250 mg/dL) 9 days after the final STZ injection. Surgery and assessments were performed by unblinded individuals, and mice were randomly assigned to experimental groups. For transplantation, mice were anesthetized with isoflurane and injected with either 5×10^6^ SC-islet cells or 4,000 IEQ cadaveric human islets underneath the kidney capsule. Transplanted SC-islet cells were derived from the AN1.1 hiPSC line either without (AN1.1 – control, n =7) or with (AN1.1 – 0.175 µM latA, n = 8) a 0.175 µM latA treatment during the first 24 hours of stage 1. Cadaveric human islets (n = 6) were purchased from Prodo Laboratories, Inc (Aliso Viejo, CA). Control mice included those that were not treated with STZ and not transplanted with any cells (n = 5) as well as mice that were treated with STZ but not transplanted with any cells (n = 7). Mice were monitored for up to 30 weeks. Mice receiving STZ but no transplanted cells were sacrificed 6 weeks after STZ treatment due to poor health. The mice transplanted with the AN1.1 – control cells were sacrificed by week 11 due to visible tumor growth. Each week, random and fasting blood glucose levels (4-6 hour fast) were recorded for each mouse (7252, Bayer).

#### In Vivo GSIS

At week 6 and 27, human C-peptide was measured in the blood of the mice after glucose injection. On the day of the assay, the mice were fasted 6 hours. Serum was then collected, followed by a 3 g/kg glucose injection (G7528, MilliporeSigma) in 0.9% sterile saline. Serum was collected again after 30 minutes. Human C-peptide from the collected serum was quantified using the Human Ultrasensitive C-Peptide ELISA kit (10-1141-01, Mercodia).

#### Glucose Tolerance Test

Glucose tolerance tests were performed in week 8 and week 28. Mice were fasted for 6 hours, and then a fasting blood glucose measurement was collected. The mice were then injected with 3 g/kg glucose in 0.9% sterile saline, and blood glucose measurements were made every 30 minutes for up to 2 hours.

#### Nephrectomy

In week 30, live nephrectomies were performed on transplanted mice to remove the kidneys containing the transplanted cells. After one week, random and fasting blood glucose measurements were made to confirm a return to a diabetic state before sacrificing the mice. The excised kidneys were processed for histological sectioning and imaging as described earlier.

### Single-cell RNA sequencing

#### Sample preparation

For experiments testing the effect of stage 1 latA treatment, HUES8 stem cells were plated at a density of 0.63 cells/ cm^2^ in 6-well plates. The differentiations were performed as normal, with the exception that 0.175 µM latA was added to half of the wells during the first 24 hours of differentiation. Differentiation batches were staggered so that all of s1d1, s1d2, s1d3, s1d4, and s2d1 cells were processed on the same day. Similarly, s3d1, s4d1, s5d1, and s6d7 cells were processed on the same day. On the specified day of differentiation, cells were dispersed from 6-well plates by incubating the cells in TrypLE for 10 minutes at 37°C and then washed with PBS. To barcode the samples so that they could be combined during sequencing, 2 million cells of each sample were resuspended in 100 µL Cell Staining Buffer (420201, BioLegend) supplemented with 10% fetal bovine serum (F2442, MilliporeSigma) for 10 minutes at 4°C. Each sample was then incubated separately in 1μg of a unique TotalSeqA antibody (394601-394611, Biolegend) in the Cell Staining Buffer for 30 minutes at 4°C. Cells were then washed twice with Cell Staining Buffer, resuspended in DMEM at 1000 cells/µL, and pooled together. Samples were submitted to the Genome Technology Access Center at McDonnel Genome Institute of Washington University in St. Louis for further processing. Briefly, samples were processed using the 10X Single Cell GEM Single Cell 3ʹ Kit v3.1 and inserted into the 10X Chromium X to generate GEM emulsions. Samples were processed according to the 10X chromium protocol CG000315 to generate single-cell libraries and verified for fragment size (4150 TapeStation System, Agilent). Samples were then sequenced using the NovaSeq 6000 (Illumina).

For experiments testing the effects of latA on stem cells in mTeSR1, HUES8 stem cells were plated at a density of 0.63 cells/ cm^2^ in 6-well plates, as normal for a differentiation. After 24 hours, either 0.175 µM latA or 1.5 µM latA in mTeSR1 was added. After an additional 24 hours, cells were collected for sequencing as described above.

#### Raw Data Processing

RStudio (version 1.3.1093) running R (version 4.0.3) and the Seurat (version 4.3.0) package were used to perform all analyses.^69^ The sequencing library was aligned and annotated with the reference human genome (hg38) from the EnsDb.Hsapeins.v86 database. Cells were demultiplexed based on enrichment of hashtag oligos (HTOs).^70^ Dead cells, doublets, and poorly sequenced cells were excluded from the data by filtering out cells containing high mitochondrial counts, exceedingly high RNA counts, and low unique and total RNA features, respectively. Thresholds for filtering out these cells can be found in Supplementary Table 2.

#### Clustering of Integrated Single-cell RNA Sequencing Datasets

Dataset integration and normalization was performed with the Seurat (version 4.3.0) package. Data containing control cells sequenced on s1d1, s1d2, s1d3, s1d4, s2d1, s3d1, s4d1, s5d1, and s6d7; and data containing latA treated cells sequenced on s1d2, s1d3, s1d4, s2d1, s3d1, s4d1, s5d1, and s6d7 were integrated based on the 2000 most variably expressed genes into a single Seurat object using *FindIntegrationAnchors* and *IntegrateData*. At this point, gene counts were adjusted using *ScaleData* and *NormalizeData*. Clustering was performed using *RunPCA*, *FindNeighbors* and *FindClusters* with 30 dimensions.

The s6d7 UMAP in Fig. 5J was obtained by integrating and clustering only s6d7 control and s6d7 latA-treated cells as described above with 30 dimensions and a resolution of 1.0. Cell types were identified by performing differential gene expression analysis using *FindAllMarkers*. The resulting gene list can be found in Supplementary Table 2. The latA-treated hESC UMAP in Supplementary Fig. 10A was obtained by clustering control hESCs, 0.175 μM latA treated hESCs, and 1.5 μM latA treated hESCs by normalizing gene counts with *ScaleData* and *NormalizeData.* Clustering was performed using *RunPCA, FindNeighbors* and *FindClusters* with 20 dimensions. All UMAPs were visualized using *RunUMAP* and *DimPlot* functions.

#### Single-cell RNA Sequencing Comparative Gene Expression Analysis

The Wilcox test method of *FindMarkers* was used to compare differential gene expression between cell types of various conditions. *VlnPlot, FeaturePlot,* and *DoHeatmap* were used to visualize differences in gene expression level. To generate volcano plots, differential gene expression analysis was performed across two conditions using *EnhancedVolcano* of the EnhancedVolcano (version 1.8.0) package. Average expression values were computed using *AverageExpression*.

### Single-nucleus multi-omic sequencing

#### Single-nuclei sample preparation

Samples were prepared using the 10X Multiome ATAC and Gene Expression (GEX) protocol (CGOOO338) to generate nuclei samples for library preparation. Briefly, cells on either s2d1 or s6d15 were single-cell dispersed by treating them with TrypLE for 10 minutes at 37°C and assessed to ensure at least 90% viability (Vi-Cell XR, Beckman Coulter). Buffers and regents were prepared fresh according to the 10X Genomics sample prep protocol (CG000505). Dispersed cells were washed with PBS containing 0.04% BSA and lysed into nuclei by treating cells with lysis buffer for 4 minutes. Nuclei samples were washed three times with wash buffer and resuspended with 10x nuclei buffer (2000207, 10X Genomics) to reach a cell concentration of approximately 5000 nuclei/µL. Prepared nuclei samples were delivered to the Genome Technology Access Center at McDonnel Genome Institute of Washington University in St. Louis for further processing. Samples were processed using the 10x Single Cell Multiome ATAC + Gene Expression v1 kit (1000283, 10X Genomics) and inserted into the 10X Chromium X to generate GEM emulsions. Samples were then processed to generate libraries according to the manufacturer’s protocol and verified for fragment size (4150 TapeStation System, Agilent). Samples were then sequenced using the NovaSeq 6000 (Illumina).

#### Analysis of single-nuclei RNA and ATAC sequencing

Raw sequencing files were processed and analyzed by mapping to the GRCh38 human reference genome using Cell Ranger Arc (version 2.0). Processed datasets were then compiled and analyzed using RStudio (version 1.3.1093) running R (version 4.0.3). The Seurat (version 4.0) and Signac (version 1.3.0) packages were primarily used for the analysis. Briefly, RNA and ATAC sequencing data files were compiled into a Seurat object files using *CreateSeuratObject* for the RNA dataset and *CreateFragmentObject* and *CreateChromatinAssay* for the ATAC dataset. Individual datasets were assessed for quality control using *NucleosomeSignal* and *TSSEnrichment*, followed with poor quality nuclei removal by excluding cells with nCount_RNA>20,000, nCount_ATAC > 20,000, nucleosome_signal > 1.50, and TSS.enrichment <1 for all conditions. ATAC Peaks were called and mapped using MACS2 by referencing the human reference genome EnsFb.Hsapeins.v86.

For comparative analysis between conditions, RNA and ATAC datasets from various conditions were integrated by setting up the integration anchor using *SelectIntegrationFeatures*, *PrepSCTIntegration*, and *FindIntegrationAnchors*. Integration was then performed using *IntegrateData* using the anchor file and used SCTransform as the normalization method. Integrated datasets were further processed using *RunTFIDF*, *RunSVD*, and *IntegrateEmbeddings* for batch correction. *FindMultiModalNeighbors*, *RunUMAP*, and *FindClusters* were then used to compute and determine clusters. Promoter accessibility information and motif enrichment were computed using *Gene-Activity* and *RunChromVAR*, respectively, of Chromvar package.

Individual clusters were computed from the integrated datasets using gene expression of generic markers. Other unidentified clusters were assessed for highly expressed genes using *FindMarkers* and the Wilcox test method. The LR test method in *FindMarkers* was used to determine enriched motif accessibility. *Coverageplot* of the Signac package was used to generate ATAC peak figures.

## Data Availability

The single-cell RNA sequencing data and single-nucleus multi-omic sequencing data from this study are available at the Gene Expression Omnibus (GEO). All other data supporting the findings of this study are available from the corresponding author upon reasonable request.

## Code Availability

Codes used for integrating and analyzing single-cell RNA sequencing and multi-omic datasets are available at https://github.com/mschmidt22 and github.com/punnaug, respectively.

## Statistics and reproducibility

Statistical analysis was performed in GraphPad Prism, version 10. The statistical tests used in each panel are listed in the figure captions. *P* values are indicated as follows: NS, not significant; \**P* < 0.05, \*\**P* < 0.01, \*\*\**P* < 0.001. All data are presented as the mean, and all error bars represent the standard error of the mean (SEM). The sample size (*n*) indicated in the figure legends are the total number of independent biological samples. For *in vitro* assays, each biological replicate was differentiated in a separate well of a tissue culture plate, either in the same batch or from multiple batches of cells. For example, the static GSIS data in Figure 2B and the dynamic GSIS data in Figure 2E came from at least two separate differentiations done at different times for each cell line, with each batch having multiple biological replicates. Results from single cell-RNA sequencing were confirmed with qRT-PCR and at the protein level with immunostaining, western blots, and colorimetric ELISA assays to measure phosphorylation state. Overall, the robustness of the major findings in this manuscript is supported by the reproducibility of these results in 4 stem cell lines over many differentiation batches.

## Author contributions

N.J.H. and J.R.M. designed all experiments. N.J.H. and S.E.G. performed all *in vitro* experiments. M.D.S., M.S., and J.R.M. performed all *in vivo* experiments. M.D.S. and P.A. performed all computational analysis. N.J.H. and J.R.M. wrote the manuscript.

All authors revised and approved the manuscript.

## Acknowledgements

This work was funded by the NIH (R01DK114233), JDRF (3-SRA-2024-1555-S-B), a Human Islet Research Network (HIRN) Catalyst Award (via NIH NIDDK U24DK104162), the Edward J. Mallinckrodt Foundation, and the Anita Palmer Corbin Trust endowed fund to J.R.M. Further support was provided by Washington University (Departments of Surgery, Medicine, and Pediatrics, including startup funds), Mid-America Transplant Services, The Foundation for Barnes-Jewish Hospital, and the Foundation for Diabetes Research to J.R.M. We thank the Genome Technology Access Center at the McDonnell Genome Institute at Washington University School of Medicine (P30CA91842) and the Washington University Diabetes Research Center (P30DK020579) for additional analysis support. N.J.H. was funded by the JDRF (1-FAC-2023-1316-A-N). This publication is solely the responsibility of the authors and does not necessarily represent the official view of NIH nor any other funder.

