## Supplemental Figures 1-21 for "Depolymerizing F-actin accelerates the exit from pluripotency to enhance stem cell-derived islet differentiation"

[1]Division of Endocrinology, Metabolism and Lipid Research
Washington University School of Medicine
MSC 8127-057-08
660 South Euclid Avenue
St. Louis, MO 63110
USA

[2]Department of Immunology
Faculty of Medicine Siriraj Hospital
Mahidol University
Bangkok 10700
Thailand

[3] Department of Biomedical Engineering
Washington University in St. Louis
1 Brookings Drive
St. Louis, MO 63130
USA

*To whom correspondence should be addressed:
Jeffrey R. Millman,

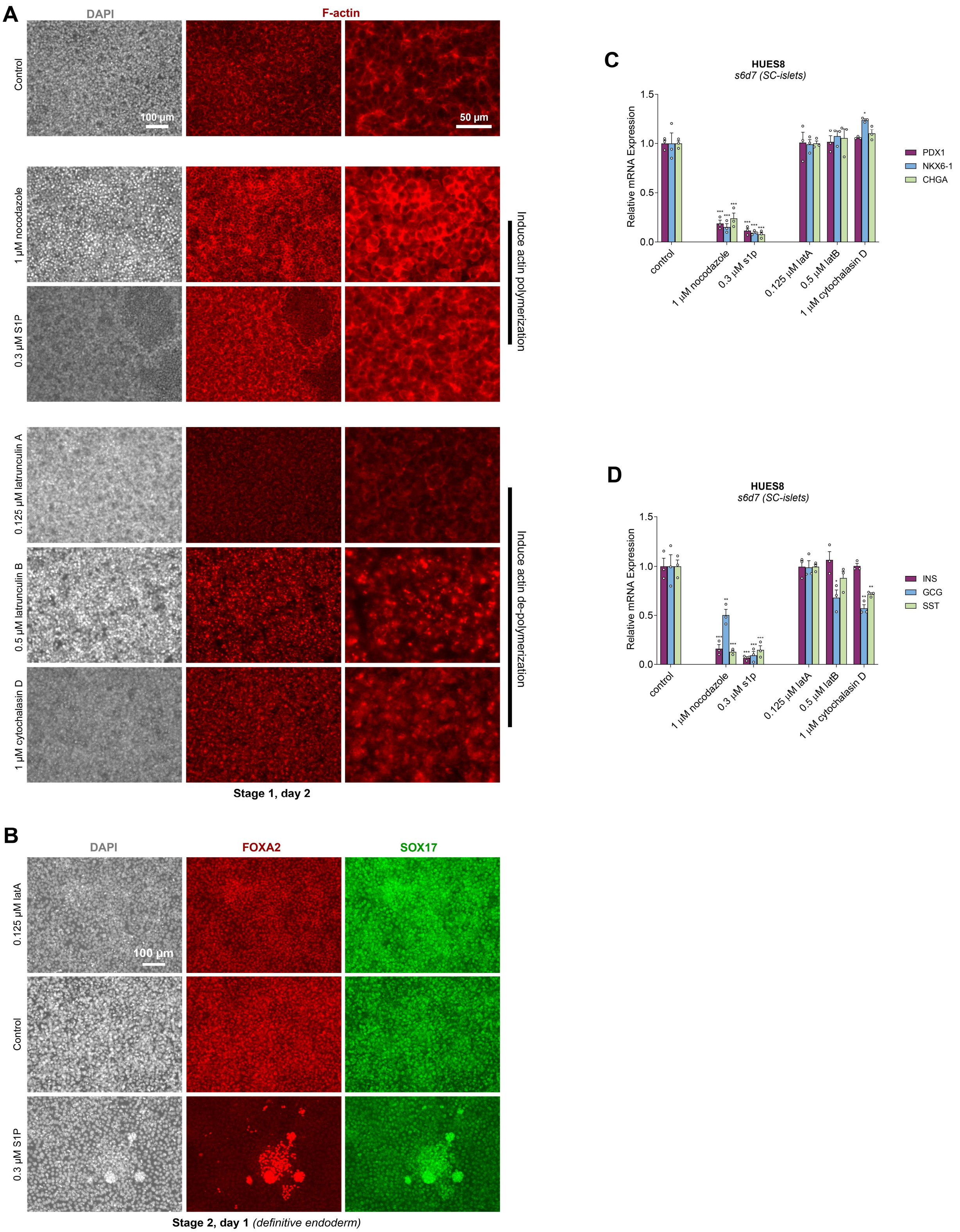
A
DAPI
F-actin
Control
100 µm
50 µm
1 µM nocodazole
0.3 µM S1P
Induce actin polymerization
0.125 µM latrunculin A
0.5 µM latrunculin B
1 µM cytochalasin D
Induce actin de-polymerization
Stage 1, day 2
B
DAPI
FOXA2
SOX17
0.125 µM latA
100 µm
Control
0.3 µM S1P
Stage 2, day 1 (definitive endoderm)
C
HUES8
s6d7 (SC-islets)
Relative mRNA Expression
1.5
1.0
0.5
0.0
control
1 µM nocodazole
0.3 µM s1p
0.125 µM latA
0.5 µM latB
1 µM cytochalasin D
PDX1
NKX6-1
CHGA
D
HUES8
s6d7 (SC-islets)
Relative mRNA Expression
1.5
1.0
0.5
0.0
control
1 µM nocodazole
0.3 µM s1p
0.125 µM latA
0.5 µM latB
1 µM cytochalasin D
INS
GCG
SST

**Supplementary Figure 1.** (**A**) Immunostaining images of F-actin on stage 1, day 2 in response to a 24-hour treatment with compounds that either induce actin polymerization (nocodazole, S1P) or that depolymerize actin (latrunculin A, latrunculin B, cytochalasin D). Scale bar = 100 µm for the two most left columns; The far-right column is a zoomed in area of the image in the center column, scale bar = 50 µm. (**B**) Immunostaining images of FOXA2 and SOX17 at the end of stage 1 in response to either latA, S1P, or no treatment for the first 24 hours of differentiation. Scale bar = 100 µm. (**C-D**) qRT-PCR of cells on stage 6, day 7 that were treated for the first 24 hours of differentiation with the indicated cytoskeletal-modulating compounds. Compounds that polymerized actin decreased expression of pancreatic endocrine genes (Dunnett's test compares each condition to the control, n = 3). All data were generated with the HUES8 cell line. All data are represented as the mean, and all error bars represent the SEM. Individual data points are shown for all bar graphs. NS, not significant; *$P < 0.05$, **$P < 0.01$, ***$P < 0.001$.

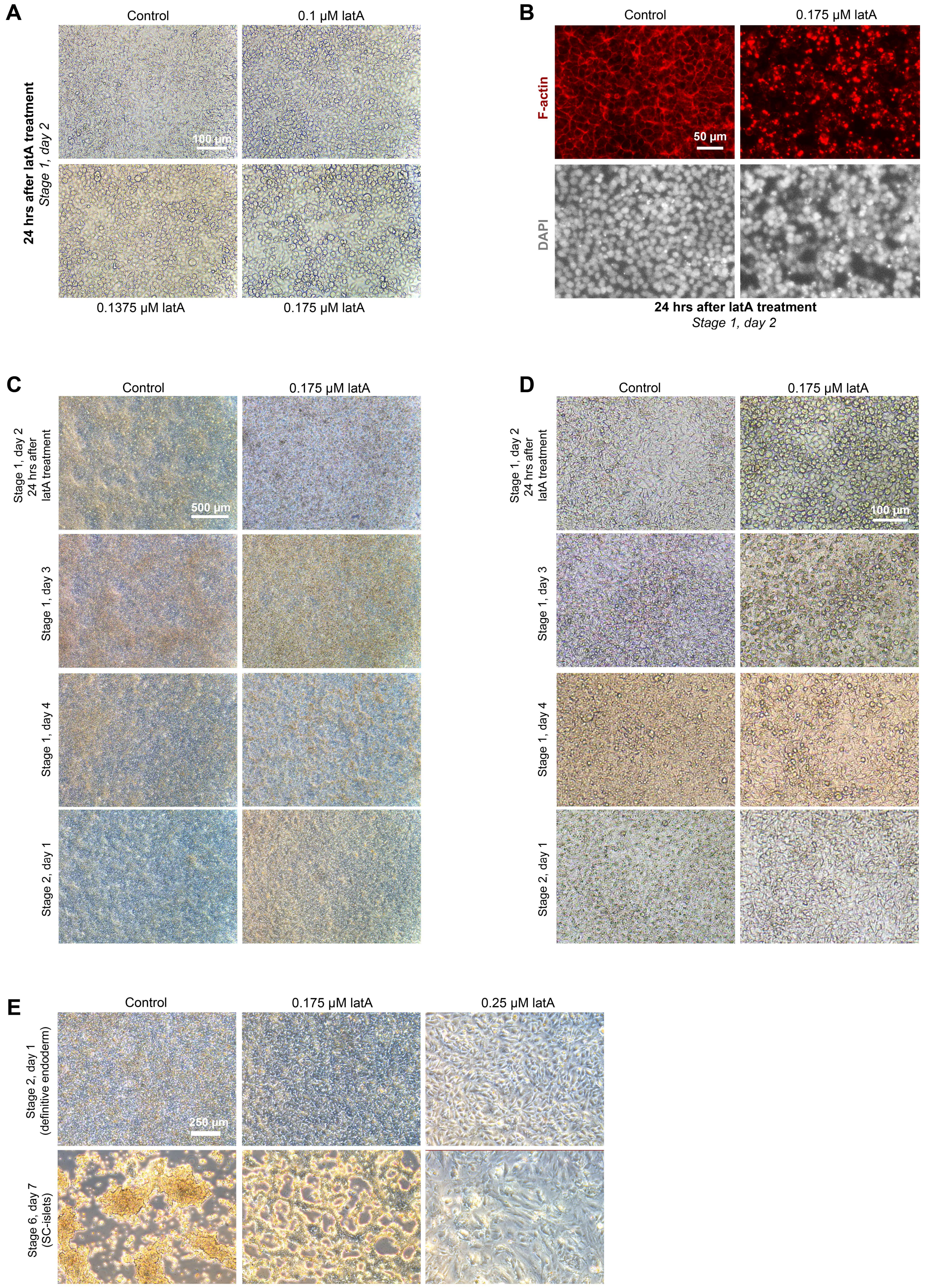
A
Control
0.1 μM latA
24 hrs after latA treatment
Stage 1, day 2
100 μm
0.1375 μM latA
0.175 μM latA
B
Control
0.175 μM latA
F-actin
50 μm
DAPI
24 hrs after latA treatment
Stage 1, day 2
C
Control
0.175 μM latA
Stage 1, day 2
24 hrs after
latA treatment
500 μm
Stage 1, day 3
Stage 1, day 4
Stage 2, day 1
D
Control
0.175 μM latA
Stage 1, day 2
24 hrs after
latA treatment
100 μm
Stage 1, day 3
Stage 1, day 4
Stage 2, day 1
E
Control
0.175 μM latA
0.25 μM latA
Stage 2, day 1
(definitive endoderm)
250 μm
Stage 6, day 7
(SC-islets)

**Supplementary Figure 2.** (**A**) Images of cells on stage 1, day 2 after a 24-hour treatment with latA (0 μM, 0.1 μM, 0.1375 μM, or 0.175 μM). Scale bar = 100 μm. (**B**) Immunostaining images of F-actin on stage 1, day 2 without and with a 24-hour treatment of 0.175 μM latA. Scale bar = 50 μm. (**C-D**) Images of cell morphology every day of stage 1 either without or with a 0.175 μM latA treatment for the first 24 hours of this stage. Notably, the latA treatment caused cells to round up, but they flattened again once latA was removed. Importantly, while increased cell death is observed throughout stage 1 with latA treatment, there should always be a confluent monolayer of cells still attached to the plate. Scale bar = 500 μm for (**C**); scale bar = 100 μm for (**D**). (**E**) Images of cell morphology on stage 2, day 1 and stage 6, day 7, demonstrating that treating the cells with too high of a latA concentration (0.25 μM) in stage 1 resulted in decreased cell density by stage 2, day 1 and non-endocrine cell types by the end of the protocol. In contrast, cells treated with an optimized latA concentration for a given cell line (0.175 μM for HUES8) produced cells with an endocrine morphology similar to the control. All images are of differentiations with the HUES8 cell line.

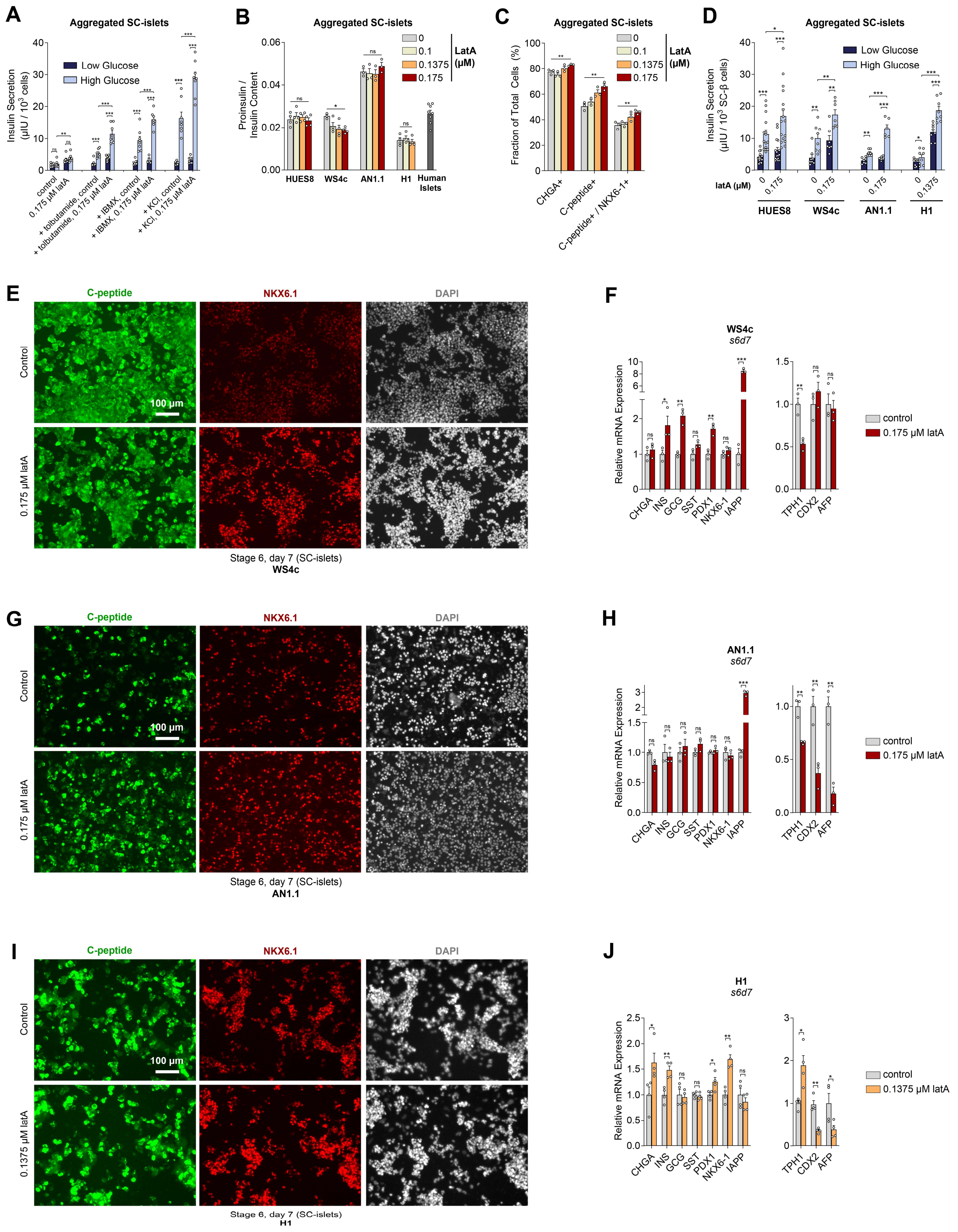
A
Aggregated SC-islets
Insulin Secretion (μIU / 10^3 cells)
Low Glucose
High Glucose
control
0.175 μM latA
+ tolbutamide, control
+ tolbutamide, 0.175 μM latA
+ IBMX, control
+ IBMX, 0.175 μM latA
+ KCl, control
+ KCl, 0.175 μM latA
B
Aggregated SC-islets
Proinsulin / Insulin Content
LatA (μM)
0
0.1
0.1375
0.175
HUES8
WS4c
AN1.1
H1
Human Islets
C
Aggregated SC-islets
Fraction of Total Cells (%)
CHGA+
C-peptide+
C-peptide+ / NKX6-1+
D
Aggregated SC-islets
Insulin Secretion (μIU / 10^3 SC-β cells)
latA (μM)
E
C-peptide
NKX6.1
DAPI
Control
0.175 μM latA
100 μm
Stage 6, day 7 (SC-islets)
WS4c
F
WS4c
s6d7
Relative mRNA Expression
CHGA
INS
GCG
SST
PDX1
NKX6-1
IAPP
TPH1
CDX2
AFP
control
0.175 μM latA
G
C-peptide
NKX6.1
DAPI
Control
0.175 μM latA
100 μm
Stage 6, day 7 (SC-islets)
AN1.1
H
AN1.1
s6d7
Relative mRNA Expression
control
0.175 μM latA
I
C-peptide
NKX6.1
DAPI
Control
0.1375 μM latA
100 μm
Stage 6, day 7 (SC-islets)
H1
J
H1
s6d7
Relative mRNA Expression
control
0.1375 μM latA

**Supplementary Figure 3.** (**A**) SC-islets generated with a 0.175 µM latA treatment during the first 24 hours of differentiation secreted more insulin in response to various secretagogues (two-way paired t-test between low and high glucose; unpaired two-way t-test between control and latA with each secretagogue at high glucose, n = 8). (**B**) Proinsulin content to insulin content ratio in stage 6 in response to latA dosing in stage 1 (one-way ANOVA, n = 4 for HUES8, WS4c, and H1, n = 3 for AN1.1, n = 8 for human islets from 2 separate donors). (**C**) Flow cytometry quantification of endocrine marker expression in stage 6 in response to increasing latA concentration in stage 1 (one-way ANOVA, n = 3). (**D**) Static GSIS of Figure 1H normalized to the number of SC-β cells rather than the total number of cells (two-way paired t-test comparing insulin secretion between low and high glucose stimulation; two-way unpaired t-test between high glucose for each cell line; n = 15-16 for HUES8, n = 6-8 for WS4c, n = 6-7 for AN1.1, n = 8 for H1, n = 11 for human islets from 2 separate donors). (**E, G, I**) Immunostaining images of stage 6, day 7 cells generated from the (**E**) WS4c, (**G**) AN1.1, and (**I**) H1 cell lines stained for C-peptide and NKX6.1. Scale bar = 100 µm. (**F, H, J**) qRT-PCR of stage 6, day 7 cells generated from the (**F**) WS4c, (**H**) AN1.1, and (**J**) H1 cell lines (two-way unpaired t-test, n = 3 for WS4c and AN1.1; n = 4 for H1). Panels (**A, C**) were performed with the HUES8 cell line. All other panels used either the HUES8, WS4c, AN1.1, or H1 cell line, as indicated. All data are represented as the mean, and all error bars represent the SEM. Individual data points are shown for all bar graphs. NS, not significant; *$P < 0.05$, **$P < 0.01$, ***$P < 0.001$.

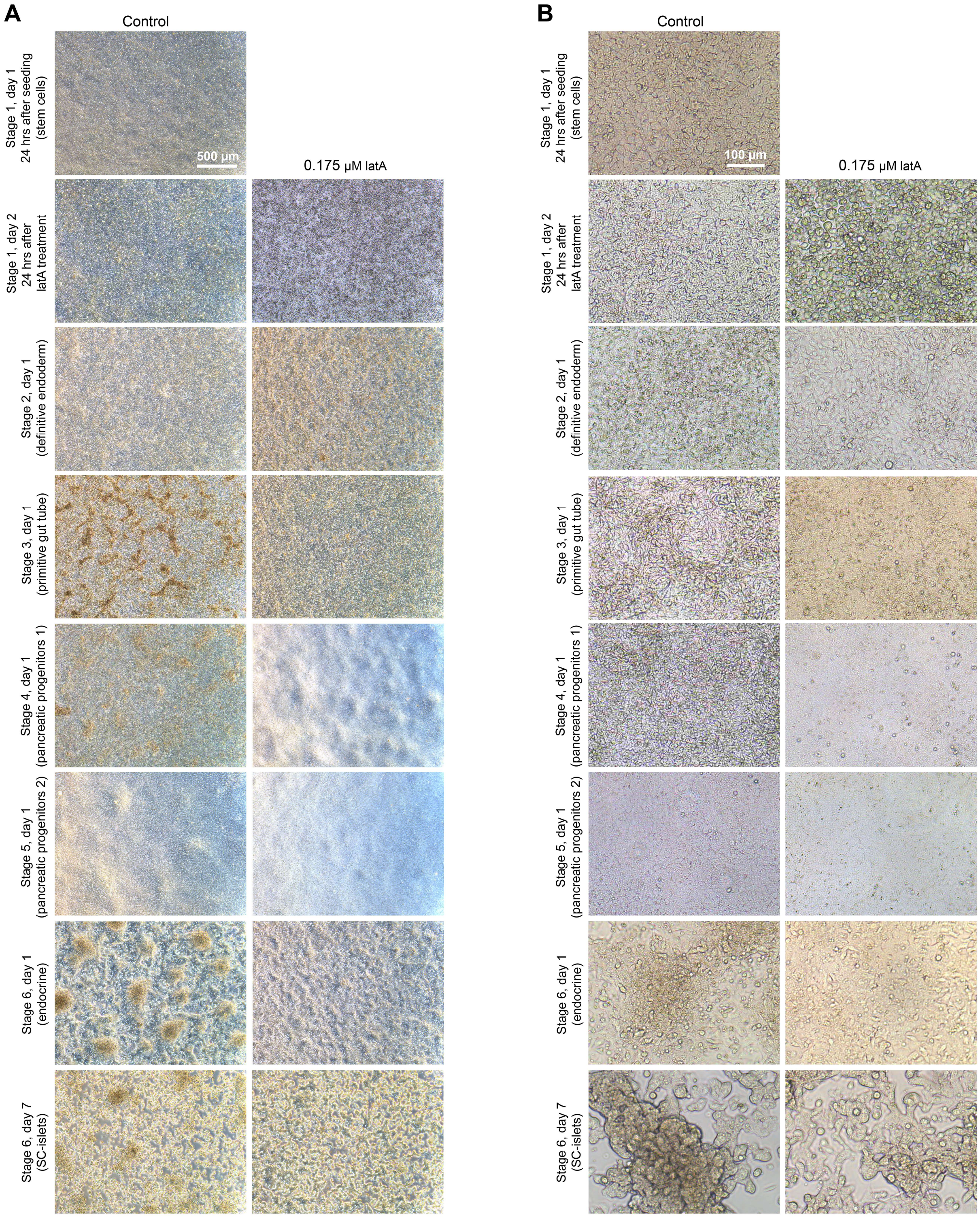


**Supplementary Figure 4.** (**A-B**) Images of the cells at each stage of the differentiation protocol to compare cell morphology between without or with a 0.175 µM latA treatment during the first 24 hours of the protocol. Notably, the latA-treated cells appeared more homogenous throughout the differentiation process. Scale bar = 500 µm for (**A**); scale bar = 100 µm for (**B**).

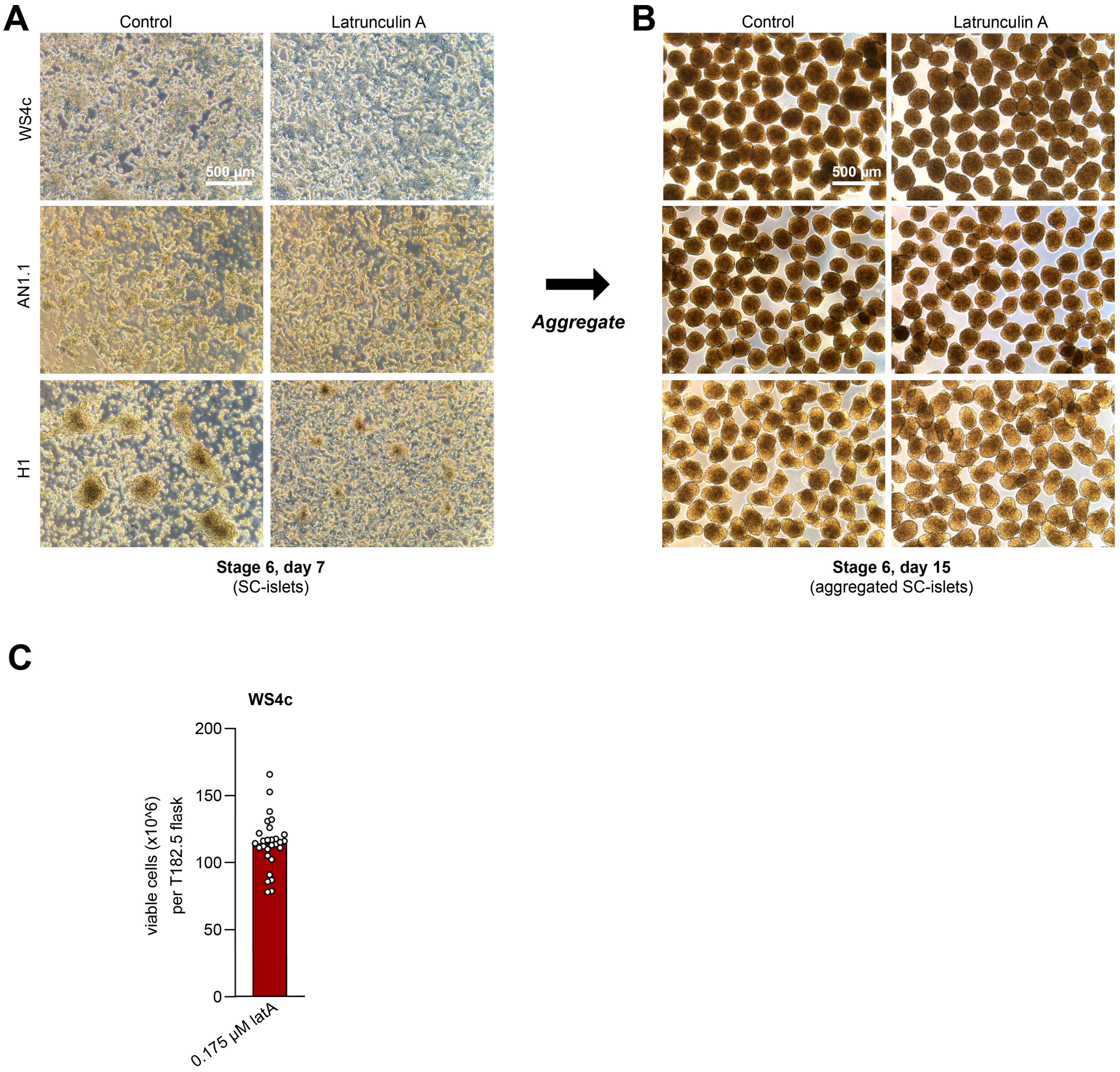


**Supplementary Figure 5.** (**A-B**) Morphology images in stage 6 of successful differentiations from the WS4c, AN1.1, and H1 cell lines both (**A**) before and (**B**) 8 days after aggregation into islet-like clusters. Differentiations were performed either without or with a latA treatment during the first 24 hours of the protocol (0.175 µM for WS4c and AN1.1, 0.1375 µM for H1). Scale bars = 500 µm. (**C**) Viable cell counts during the aggregation step on stage 6, day 7 for differentiations performed with the WS4c cell line with a 0.175 µM latA treatment in culture flasks with a surface area of 182.5 cm$^2$.

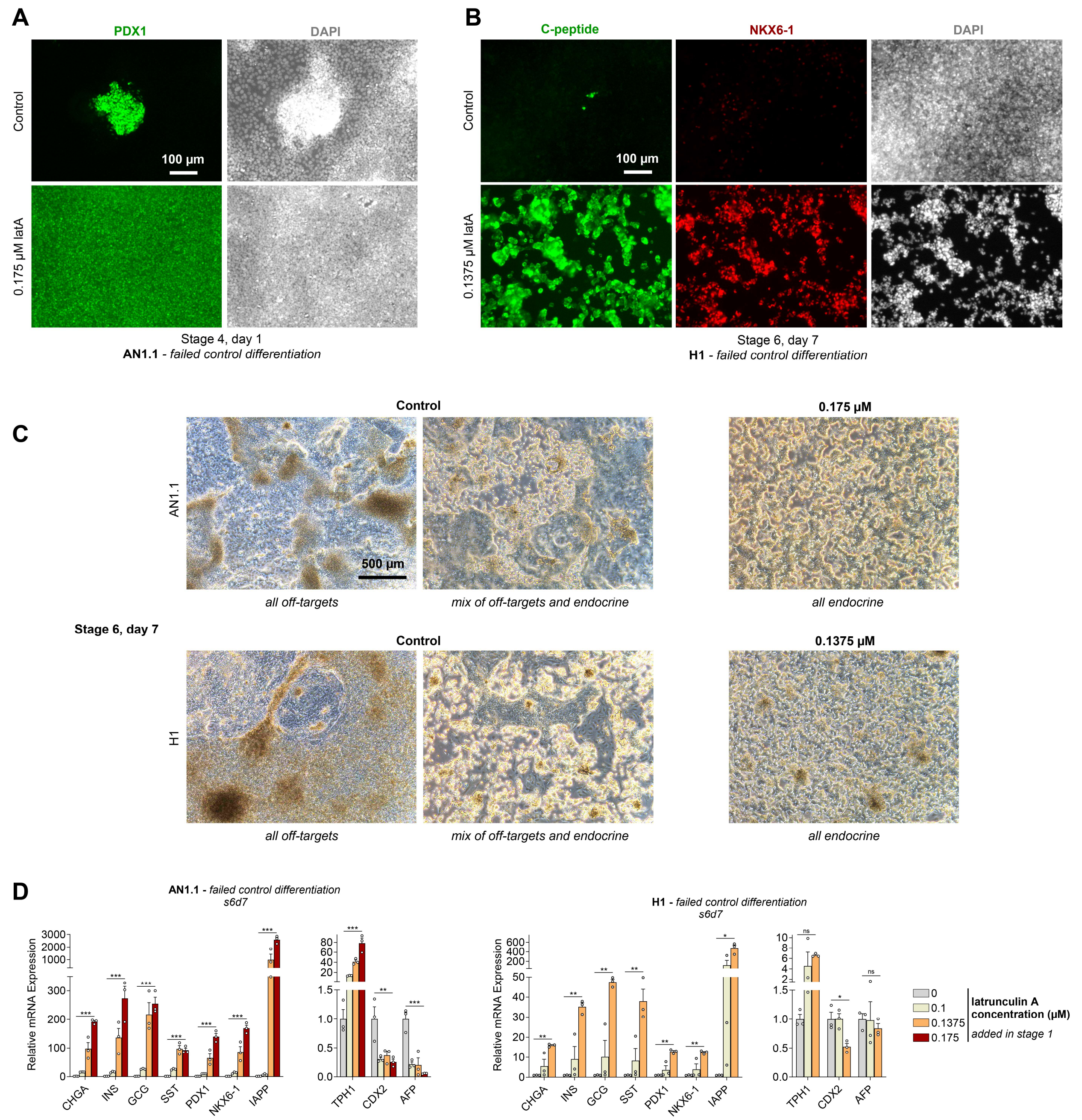


**Supplementary Figure 6.** (**A**) Immunostaining images of stage 4, day 1 cells generated from the AN1.1 line stained for PDX1. These images show a batch of cells that failed to make pancreatic progenitors with the normal differentiation protocol, while treating this same batch of cells with 0.175 µM latA during stage 1 rescued induction of PDX1+ pancreatic progenitors. Scale bar = 100 µm. (**B**) Immunostaining images of stage 6, day 7 cells generated from the H1 line stained for C-peptide and NKX6.1. These images show a batch of cells that failed to make endocrine cells with the normal differentiation protocol, while treating this same batch of cells with 0.1375 µM latA during stage 1 rescued SC-β cell generation. Scale bar = 100 µm. (**C**) Morphology images of cells generated with the AN1.1 and H1 cell lines on stage 6, day 7. The control differentiations for these cell lines would often generate non-endocrine cell types (*left*), while treating the cells with latA during the first 24 hours of stage 1 would rescue endocrine formation in these same batches of cells (*right*). (**D**) qRT-PCR of stage 6, day 7 cells differentiated from the AN1.1 and H1 cell lines from batches of cells in which the control differentiation failed. Increasing latA concentration in stage 1 progressively increased expression of pancreatic endocrine genes by stage 6, day 7 (one-way ANOVA for each gene, n = 3). All data are represented as the mean, and all error bars represent the SEM. Individual data points are shown for all bar graphs. NS, not significant; $^*P < 0.05$, $^{**}P < 0.01$, $^{***}P < 0.001$.

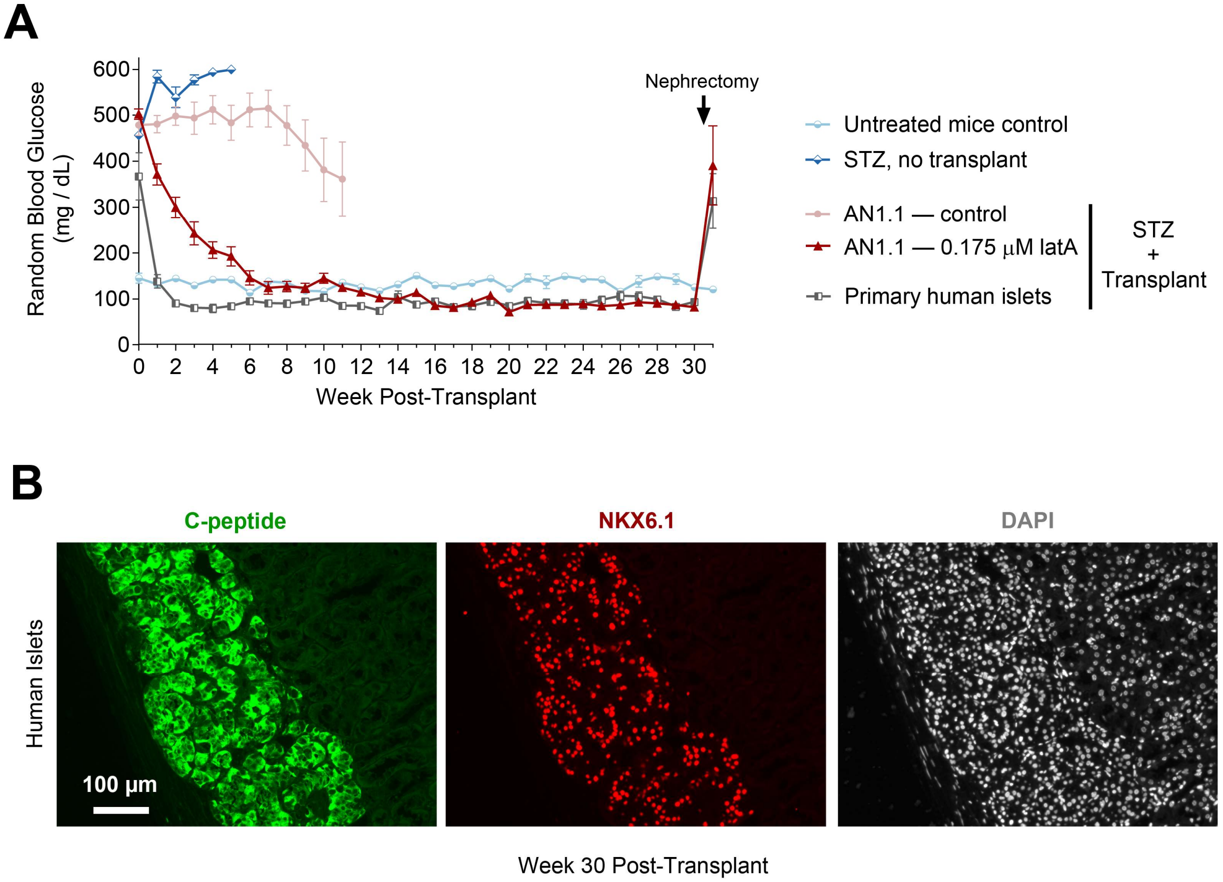


**Supplementary Figure 7.** (**A**) Random blood glucose measurements of diabetic mice that had cells transplanted underneath the kidney capsule using either primary human islets or SC-islets generated with the AN1.1 stem cell line. These SC-islets were produced either without or with the 0.175 µM latA treatment during stage 1 (untreated control: n = 5 mice; STZ, no transplant: n = 7 mice; AN1.1 – control: n = 7 mice; AN1.1 – 0.175 µM latA: n = 8 mice; primary human islets: n = 6 mice). (**B**) Immunostaining of histology sections taken from mouse kidneys transplanted with primary human islets, showing the presence of human β cells (C-peptide+/NKX6.1+) 30 weeks after transplantation. Scale bar = 100 µm. (**C**) Measurement of circulating levels of mouse C-peptide, demonstrating that mice receiving STZ and the SC-islet transplant had low levels of mouse C-peptide compared to the untreated control mice (two-way unpaired t-test). All data are represented as the mean, and all error bars represent the SEM. Individual data points are shown for all bar graphs. NS, not significant; *$P < 0.05$, **$P < 0.01$, ***$P < 0.001$.

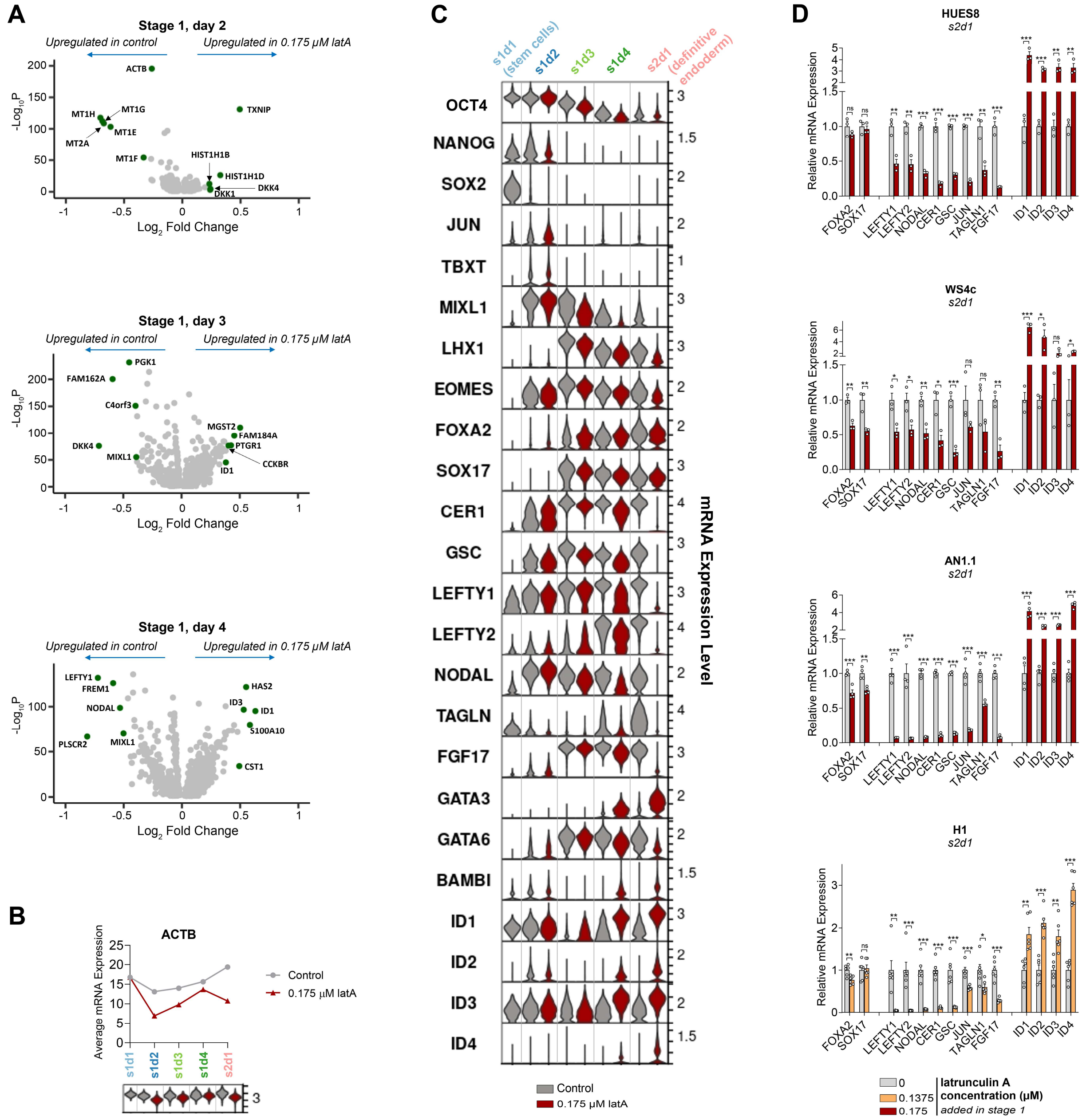


**Supplementary Figure 8.** (**A**) Volcano plots from single cell RNA-sequencing of genes differentially expressed throughout stage 1 with latA treatment. (**B**) Average mRNA expression of the whole cell population and the corresponding violin plots for ACTB at each day of stage 1. (**C**) Violin plots from single cell RNA-sequencing of genes differentially expressed throughout stage 1 with latA treatment. Panels (**A-C**) correspond to the single cell-RNA sequencing data presented in Figure 4. All sequencing data was performed with HUES8. (**D**) qRT-PCR of genes differentially expressed at the end of stage 1, demonstrating similar trends in the expression of these specific genes in response to latA treatment in all 4 cell lines (two-way unpaired t-test; n = 3 for HUES8 and WS4c, n = 4 for AN1.1, n = 6 for H1). All data are represented as the mean, and all error bars represent the SEM. Individual data points are shown for all bar graphs. NS, not significant; *$P < 0.05$, **$P < 0.01$, ***$P < 0.001$.

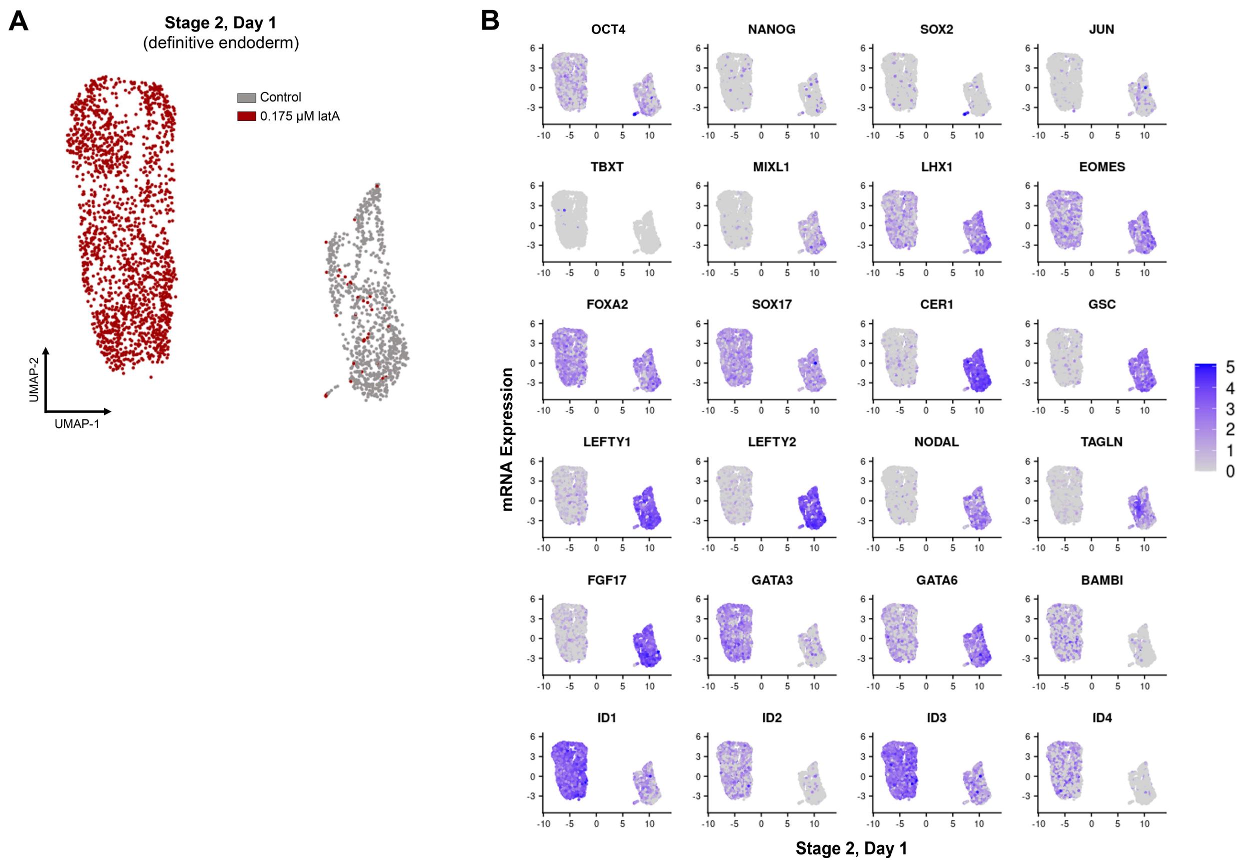


**Supplementary Figure 9.** (**A**) Single cell RNA-sequencing UMAP on stage 2, day 1 of cells treated without and with 0.175 μM latA for the first 24 hours of differentiation. (**B**) Feature plots for select genes on stage 2, day 1. All sequencing data was performed with HUES8 and corresponds to the single cell-RNA sequencing data presented in Figure 4.

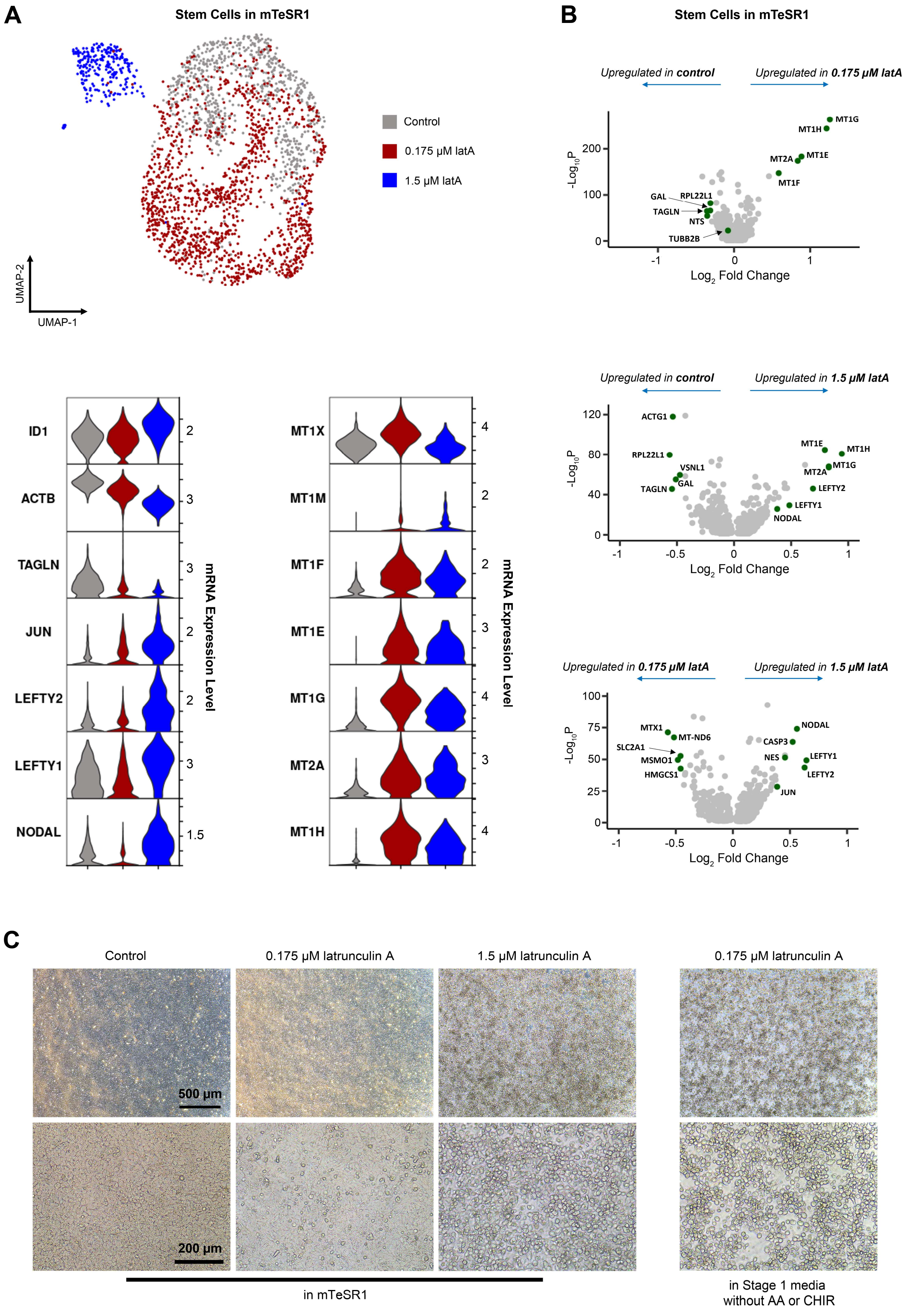
A
Stem Cells in mTeSR1
Control
0.175 μM latA
1.5 μM latA
UMAP-2
UMAP-1
B
Stem Cells in mTeSR1
Upregulated in control
Upregulated in 0.175 μM latA
Upregulated in 1.5 μM latA
Upregulated in 0.175 μM latA
Upregulated in 1.5 μM latA
Log2 Fold Change
-Log10P
ID1
ACTB
TAGLN
JUN
LEFTY2
LEFTY1
NODAL
MT1X
MT1M
MT1F
MT1E
MT1G
MT2A
MT1H
mRNA Expression Level
C
Control
0.175 μM latrunculin A
1.5 μM latrunculin A
0.175 μM latrunculin A
500 μm
200 μm
in mTeSR1
in Stage 1 media
without AA or CHIR

**Supplementary Figure 10.** (**A**) Single cell RNA-sequencing UMAP and violin plots of HUES8 stem cells treated with either 0 μM, 0.175 μM, or 1.5 μM latA for 24 hours in mTeSR1. (**B**) Volcano plots showing genes differentially expressed between these latA treatments. (**C**) Morphology images of stem cells treated with either 0 μM, 0.175 μM, or 1.5 μM latA for 24 hours in mTeSR1. A much higher latA concentration (1.5 μM) was required to induce a round morphology in mTeSR1 when compared to cells cultured in stage 1 media (0.175 μM), regardless of the presence of AA and CHIR. Scale bars = 500 μm (*top row*) and 200 μM (*bottom row*).

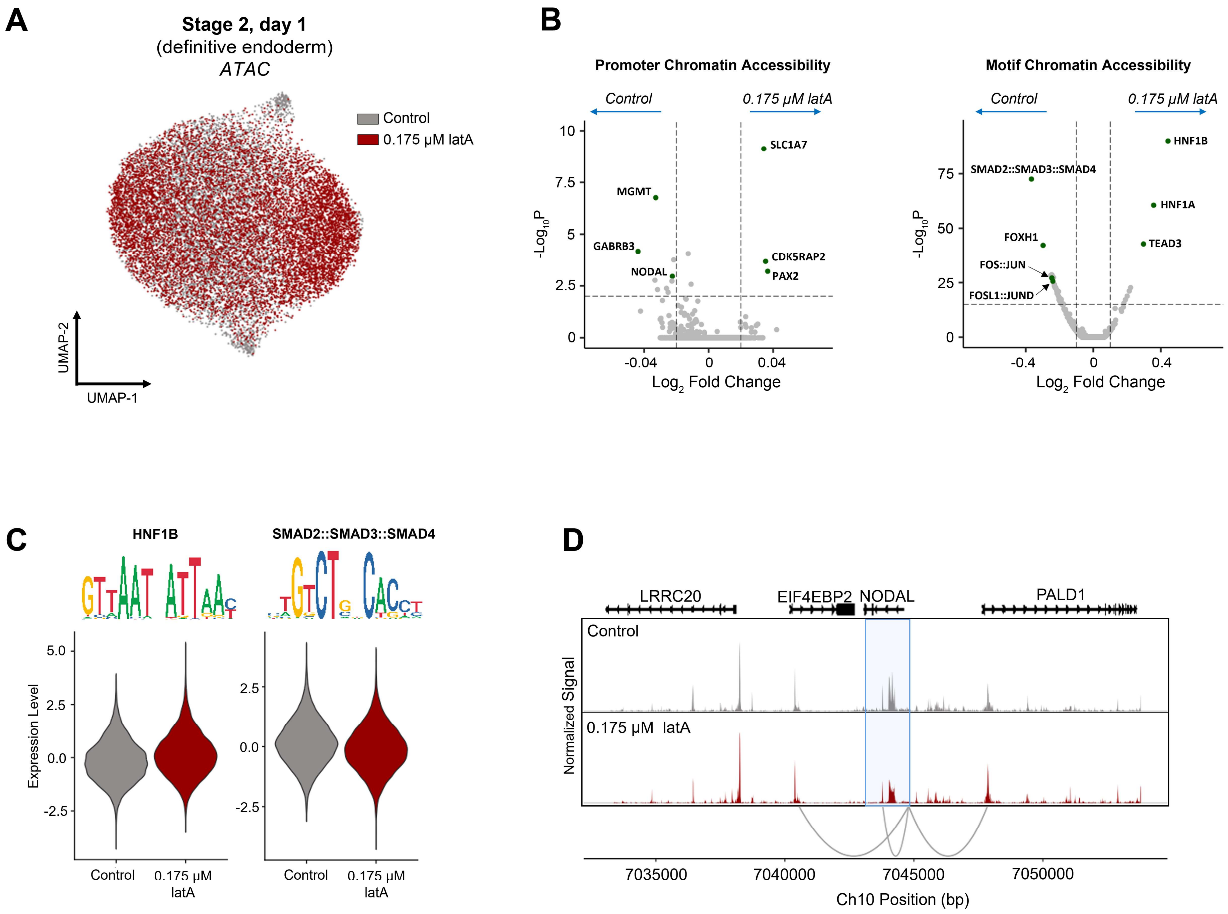


**Supplementary Figure 11.** (**A**) Single nuclei ATAC-sequencing UAMP at the end of stage 1 of HUES8 cells treated without and with 0.175 µM latA for the first 24 hours of differentiation. (**B**) Volcano plots showing differences in chromatin accessibility for gene promoters and the chromatin accessibility of transcription factor binding motifs at the end of stage 1. (**C**) Motif enrichment analysis in latA-treated cells at the end of stage 1. (**D**) ATAC plot showing the chromatin accessibility around the NODAL genomic region.

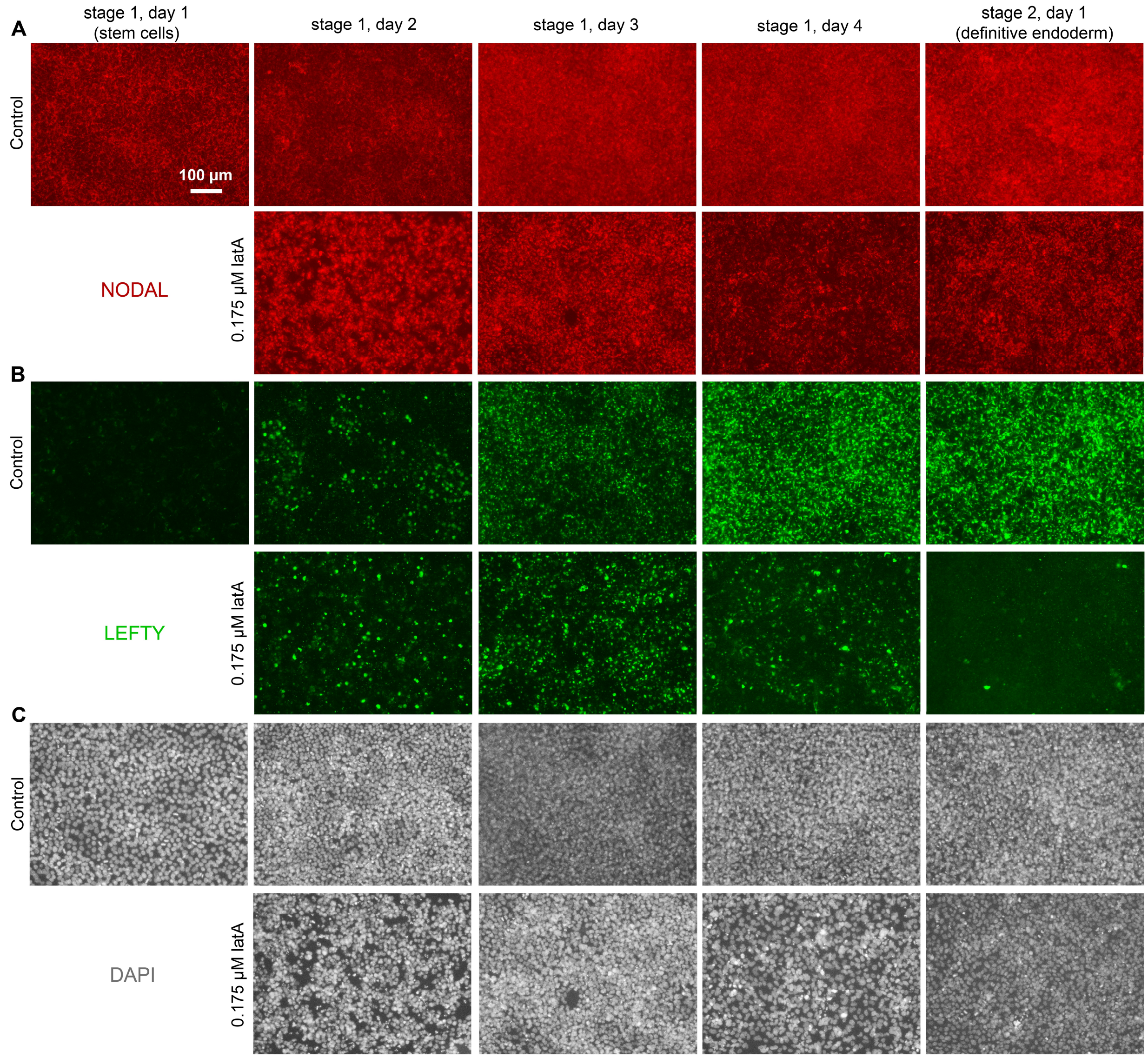


**Supplementary Figure 12.** Immunostaining images of cells each day of stage 1 treated without or with 0.175 μM latA for the first 24 hours of differentiation and stained for (**A**) NODAL, (**B**) LEFTY 1/2, and (**C**) DAPI. The images on stage 2, day 1 shown here are the same as in Fig. 4D to show comparison to the other days in stage 1. Scale bar = 100 μm.

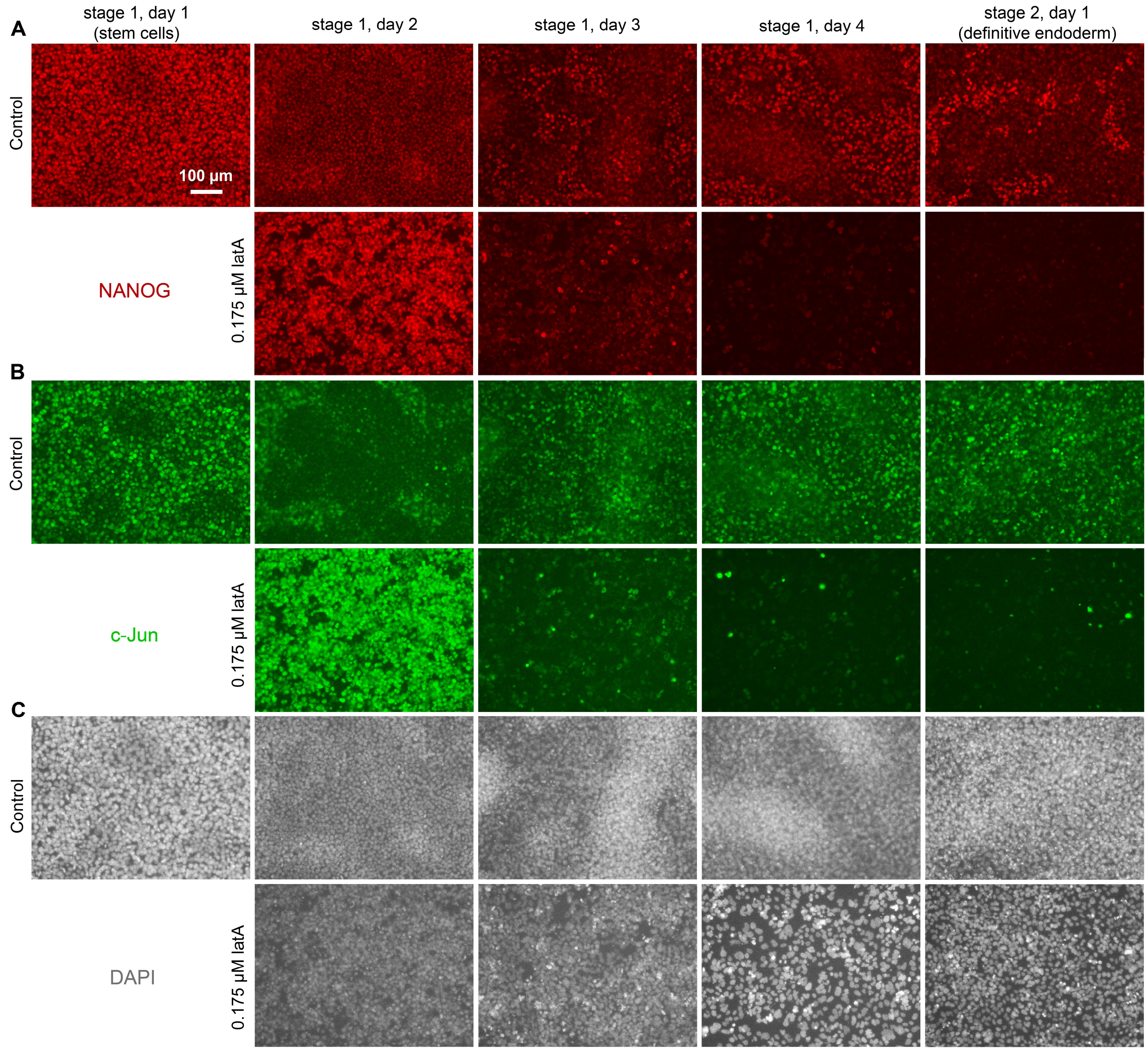


**Supplementary Figure 13.** Immunostaining images of cells each day of stage 1 treated without or with 0.175 µM latA for the first 24 hours of differentiation and stained for (**A**) NANOG, (**B**) c-Jun, and (**C**) DAPI. The images on stage 2, day 1 shown here are the same as in Fig. 4D to show comparison to the other days in stage 1. Scale bar = 100 µm.

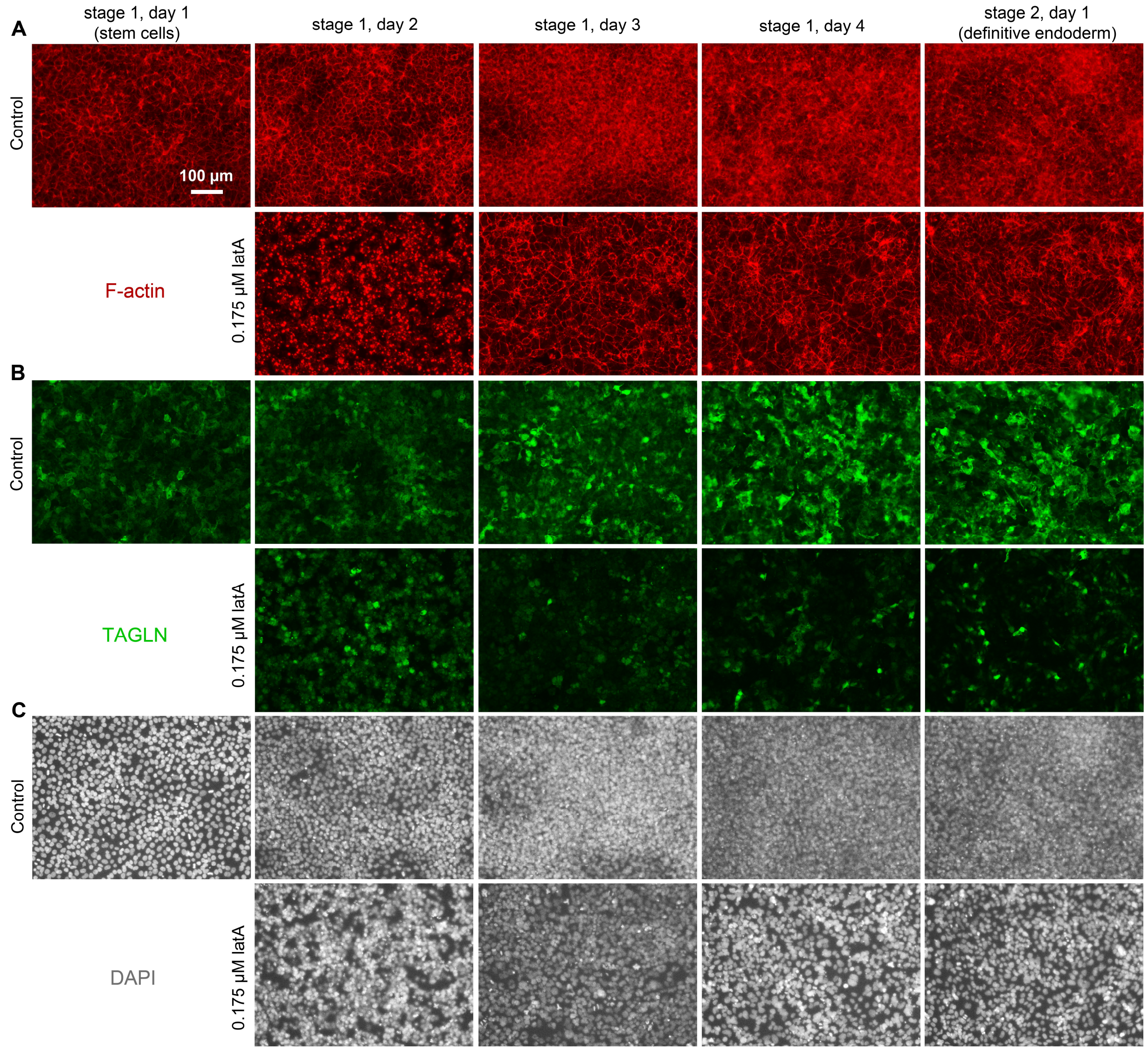


**Supplementary Figure 14.** Immunostaining images of cells each day of stage 1 treated without or with 0.175 µM latA for the first 24 hours of differentiation and stained for (**A**) F-actin, (**B**) TAGLN, and (**C**) DAPI. The images on stage 2, day 1 shown here are the same as in Fig. 4D to show comparison to the other days in stage 1. Scale bar = 100 µm.

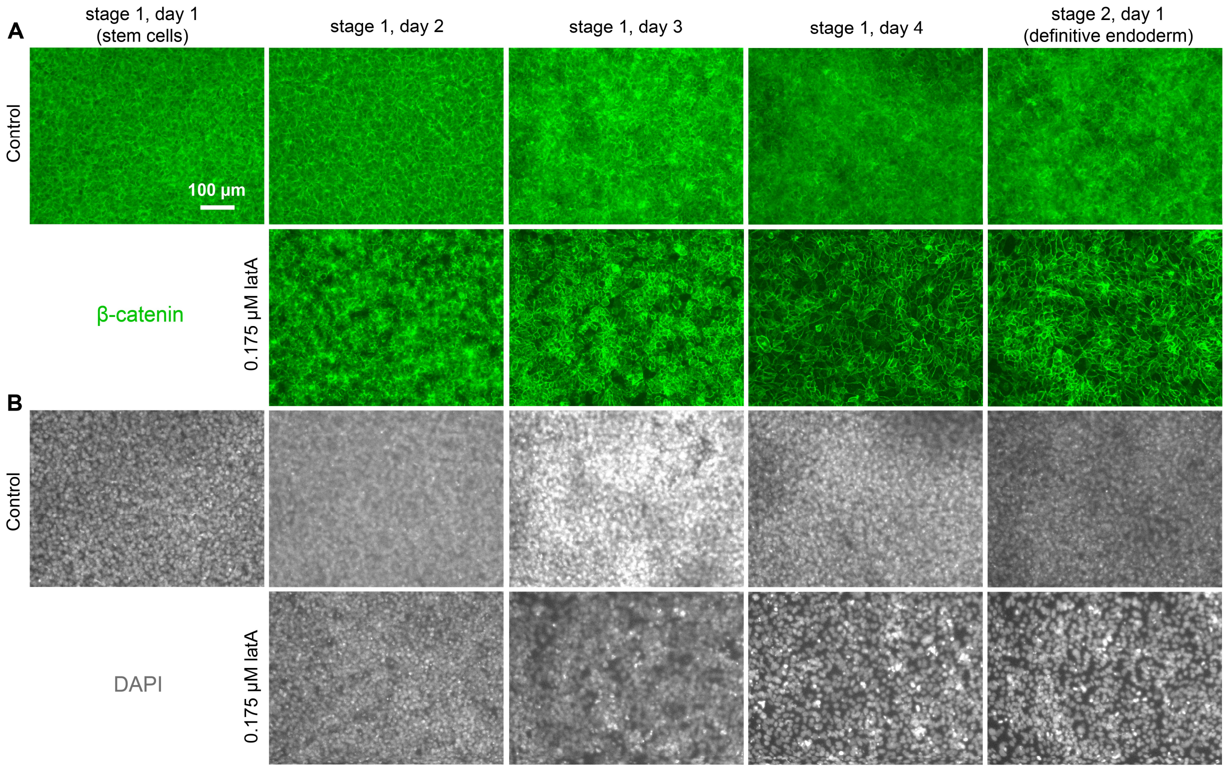


**Supplementary Figure 15.** Immunostaining images of cells each day of stage 1 treated without or with 0.175 μM latA for the first 24 hours of differentiation and stained for (**A**) β-catenin and (**B**) DAPI. Scale bar = 100 μm.

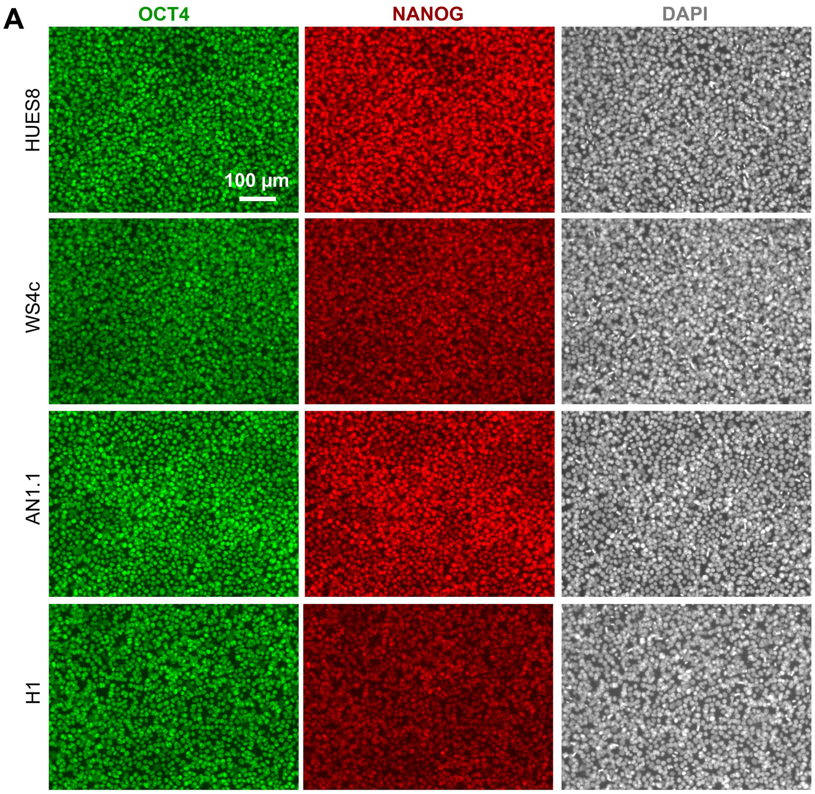


**stage 1, day 1**
(stem cells)

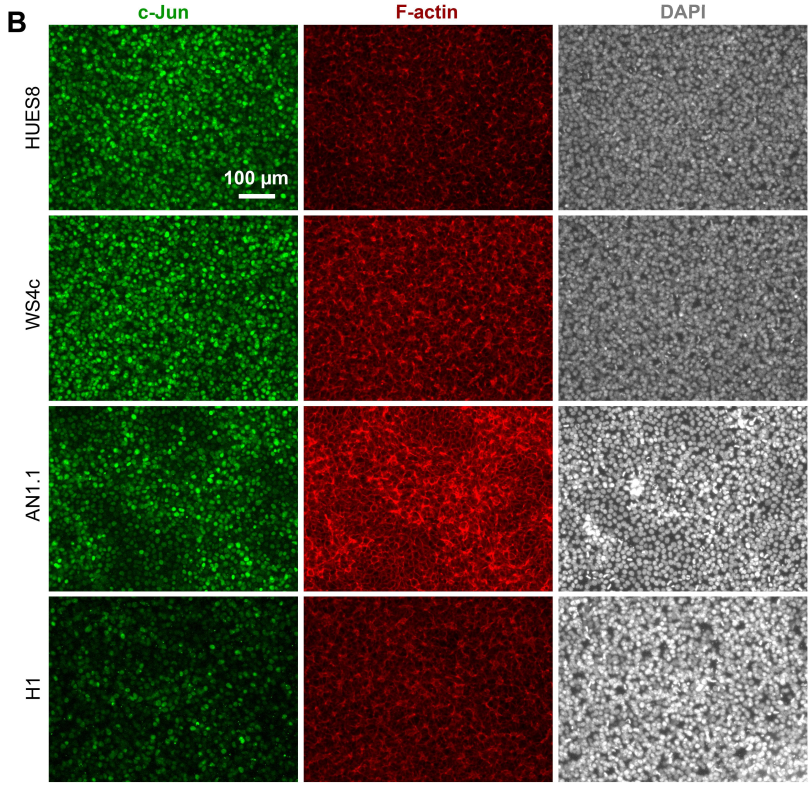


**stage 1, day 1**
(stem cells)

**Supplementary Figure 16.** (**A**) Immunostaining images of the HUES8, WS4c, AN1.1, and H1 hPSC lines stained for the pluripotency markers OCT4 and NANOG. Scale bar = 100 μm. (**B**) Immunostaining images of the HUES8, WS4c, AN1.1, and H1 hPSC lines stained for c-Jun and F-actin. Scale bar = 100 μm.

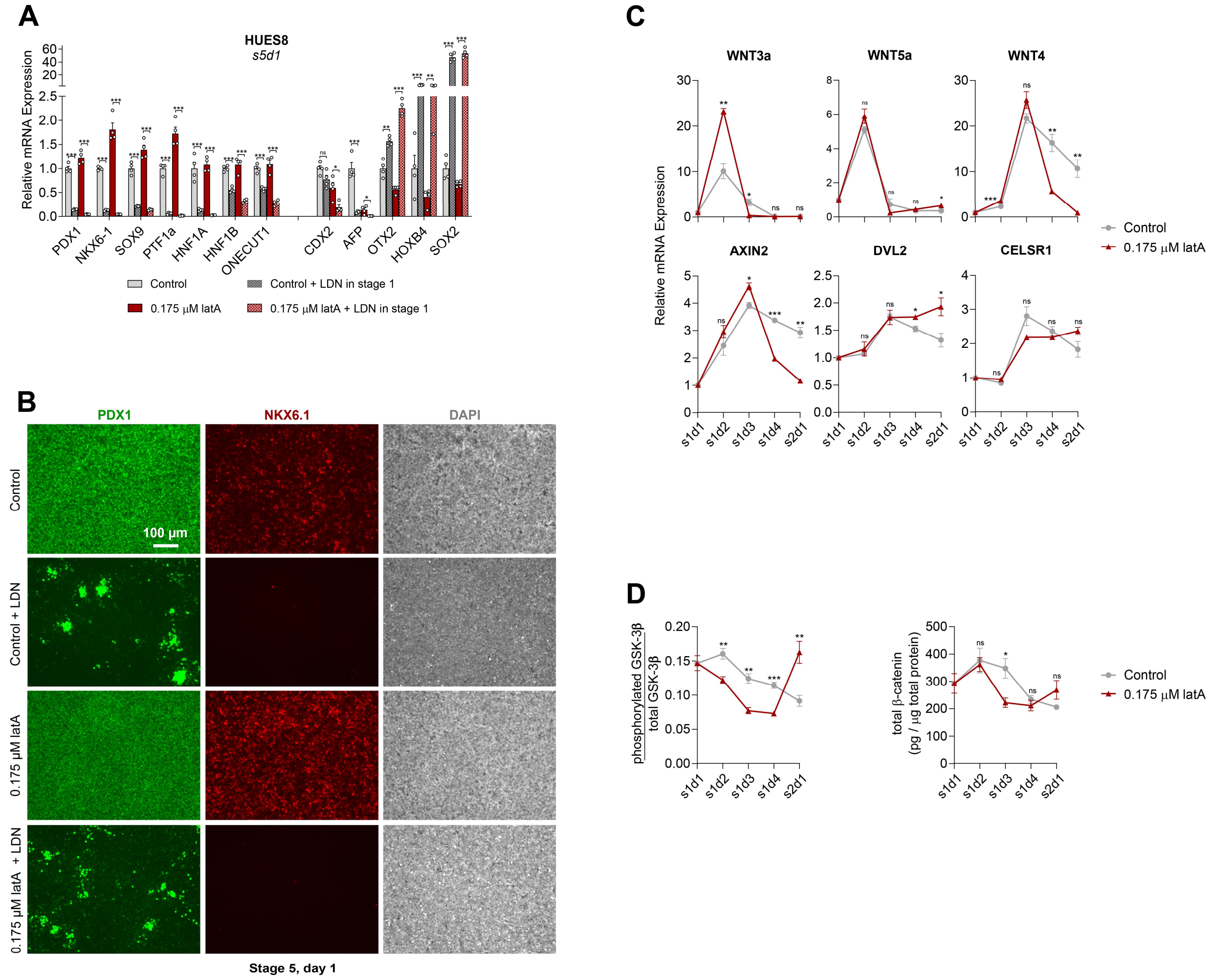


**Supplementary Figure 17.** (**A**) qRT-PCR of cells on stage 5, day 1 for pancreatic markers (*left side*) and markers of other endodermal lineages (*right side*). Cells were treated without or with 0.175 μM latA for the first 24 hours of differentiation. The BMP inhibitor LDN193189 was also added to the specified conditions starting on stage 1, day 2 through the end of stage 1 (two-way unpaired t-test for each gene, n = 4). (**B**) Immunostaining images of the conditions specified in (**A**) on stage 5, day 1 stained for PDX1 and NKX6.1. (**C**) qRT-PCR each day of stage 1 of genes associated with WNT signaling in cells treated without or with 0.175 μM latA. (**D**) ELISAs measuring the ratio of phosphorylated to total GSK-3β protein as well as total β-catenin protein every day of stage 1 without or with latA treatment (two-way unpaired t-test at each time point, n = 4). (**E**) Western blots on stage 5, day 1 for NKX6.1 (n = 3). The blots for NKX6-1 were stripped and re-probed for GAPDH, which was used as the loading control. Unprocessed western blot scans are shown in Supplementary Figure 21. All assays were performed with the HUES8 cell line. All data are represented as the mean, and all error bars represent the SEM. Individual data points are shown for all bar graphs. NS, not significant; *$P < 0.05$, **$P < 0.01$, ***$P < 0.001$.

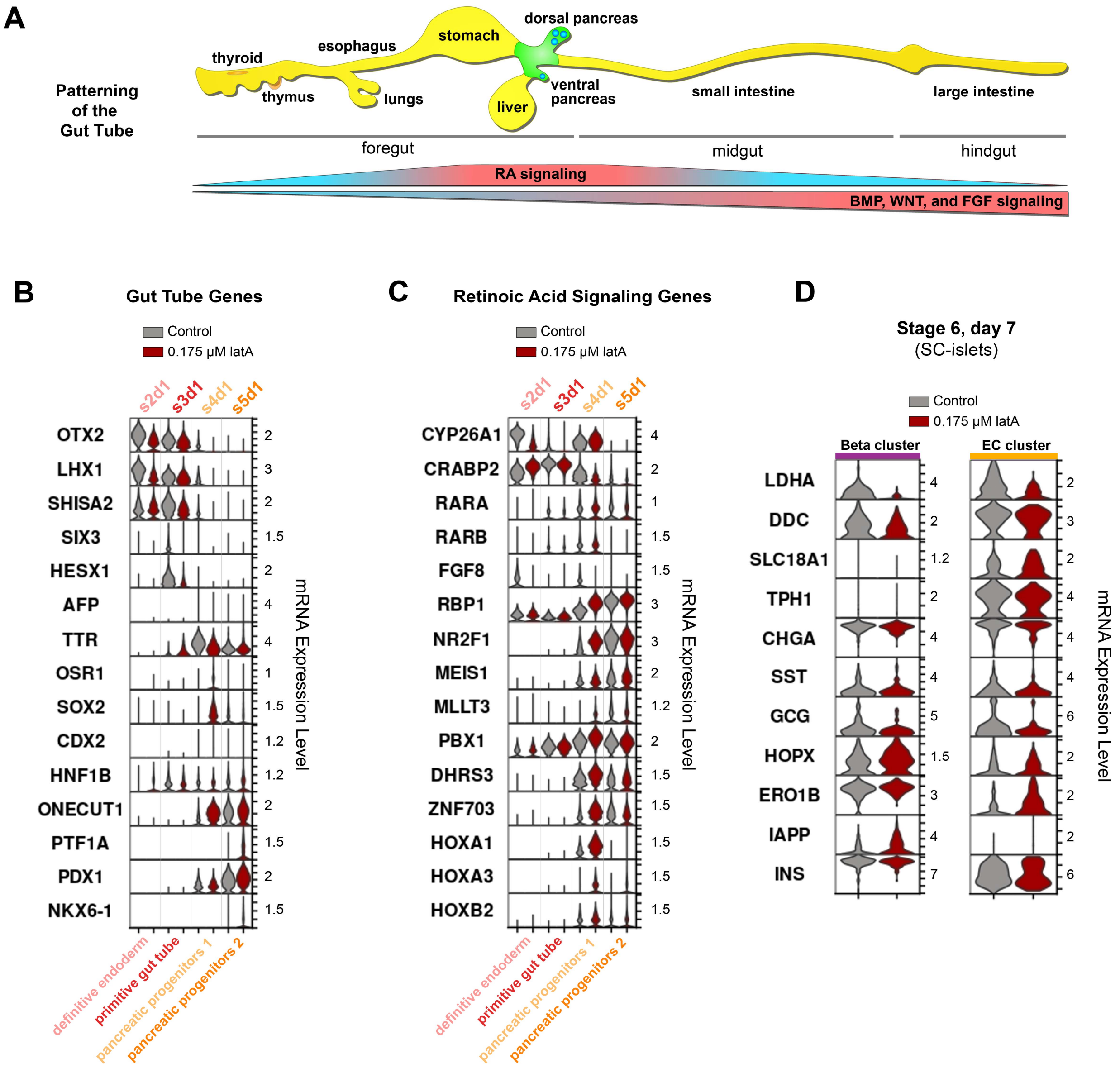


**Supplementary Figure 18.** (**A**) Gradients of signaling factors pattern the gut tube into different organ-forming regions during development. The SC-islet differentiation protocol shown in Figure 2A attempts to replicate these signaling dynamics to progressively specify the gut tube (*yellow*), then the pancreatic region (*green*), and finally the endocrine portion of the pancreas (*blue*). (**B-C**) Violin plots from the single cell RNA-sequencing data presented in Figure 5 at the intermediate stages of differentiation (s2d1, s3d1, s4d1, and s5d1) either without or with latA treatment. (**D**) Violin plots from the beta cell and enterochromaffin cell clusters on stage 6, day 7 from the single cell RNA-sequencing data presented in Figure 5.

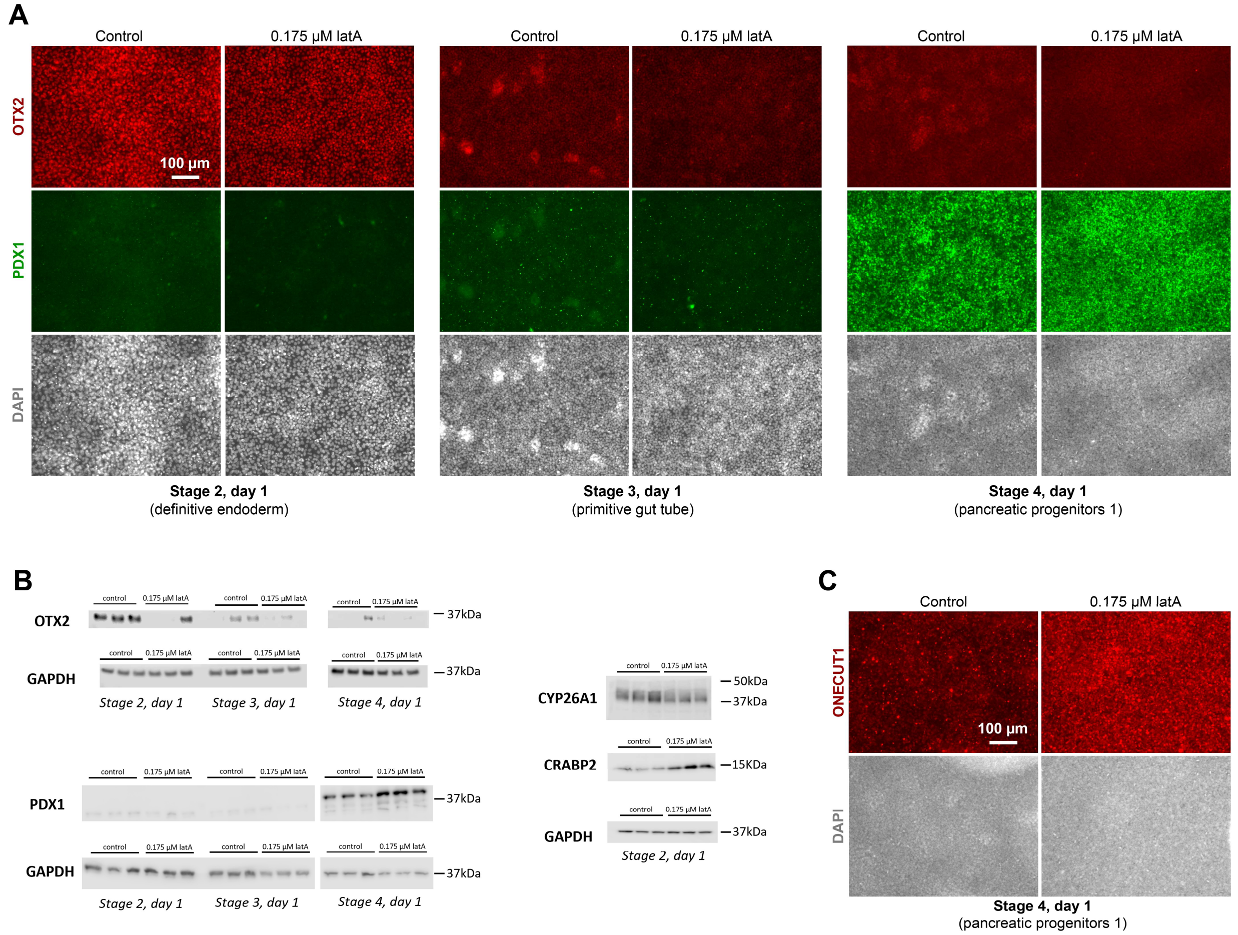


**Supplementary Figure 19.** (**A**) Immunostaining images of cells either without or with latA treatment at the intermediate stages of differentiation (s2d1, s3d1, and s4d1) stained for OTX2 and PDX1. Scale bar = 100 μm. (**B**) Western blots for markers in cells either without or with latA treatment at the intermediate stages of differentiation (s2d1, s3d1, and s4d1) (n = 3). Blots for each marker were stripped and re-probed for GAPDH, which was used as the loading control. Unprocessed western blot scans are shown in Supplementary Figure 21. (**C**) Immunostaining images of cells either without or with latA treatment at stage 4, day 1 stained for ONECUT1. Scale bar = 100 μm. All assays were performed with the HUES8 cell line.

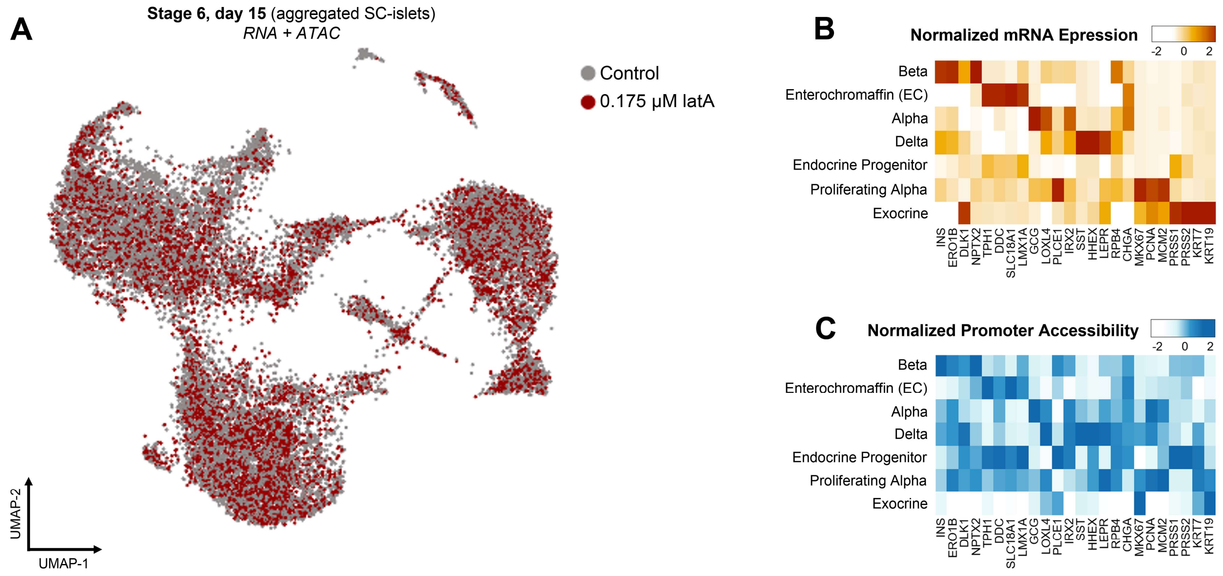


**Supplementary Figure 20.** (**A**) Single-nuclei multi-omic sequencing UMAP on stage 6, day 15 combing both RNA expression and ATAC chromatin accessibility for each cell. (**B-C**) Heatmaps of both the (**B**) mRNA expression and (**C**) chromatin accessibility for these gene promoters in each cluster.

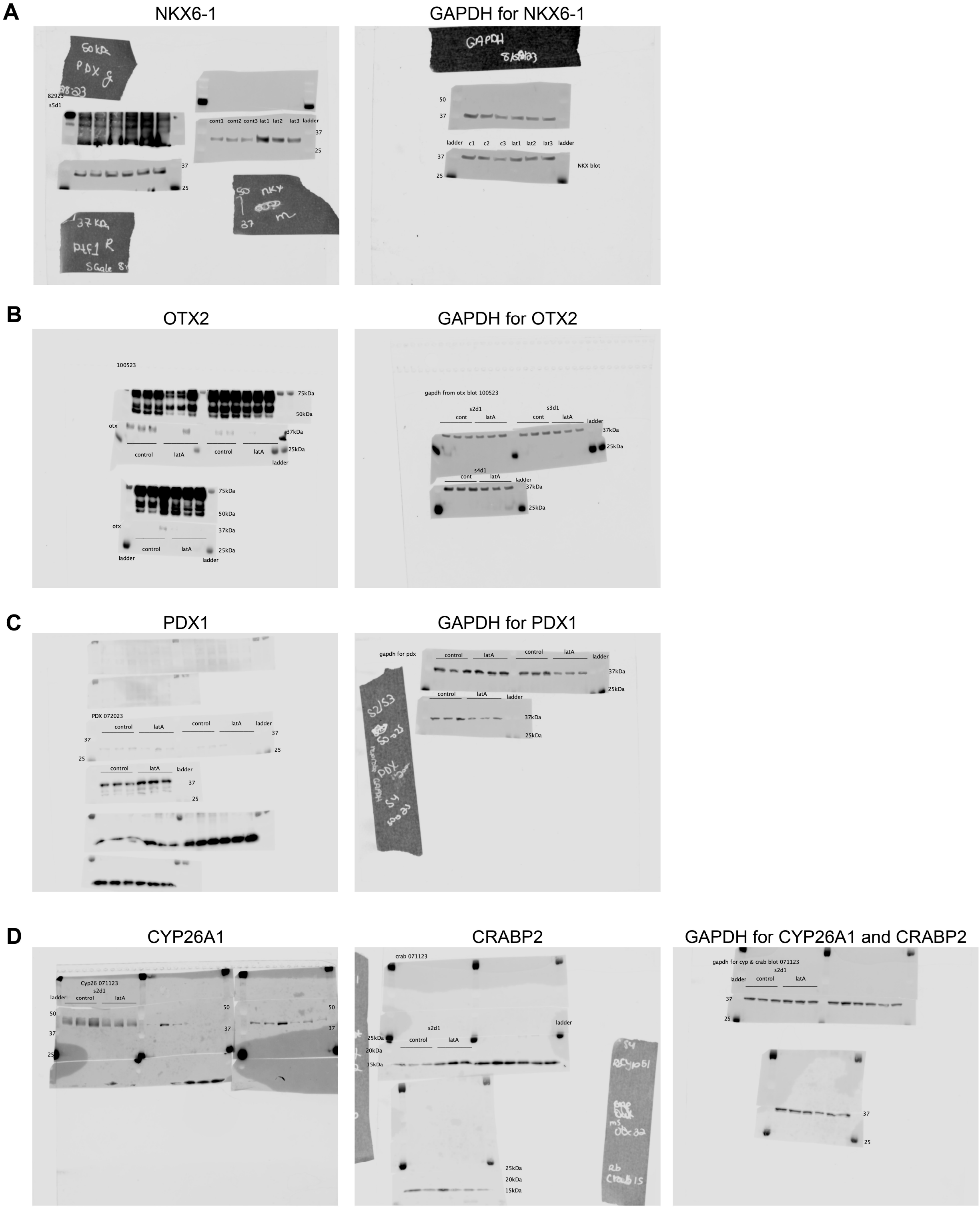


**Supplementary Figure 21.** Unprocessed scans of western blots for (**A**) NKX6-1 on s5d1, (**B**) OTX2 on s2d1, s3d1, and s4d1, (**C**) PDX1 on s2d1, s3d1, and s4d1, and (**D**) CYP26A1 and CRABP2 on s2d1. Blots for each marker were stripped and re-probed for GAPDH, which was used as the loading control. The relevant blots in each image are labeled.
